# Extracellular vesicles of Euryarchaeida: precursor to eukaryotic membrane trafficking

**DOI:** 10.1101/2023.03.03.530948

**Authors:** Joshua Mills, L. Johanna Gebhard, Florence Schubotz, Anna Shevchenko, Daan R. Speth, Yan Liao, Iain G. Duggin, Anita Marchfelder, Susanne Erdmann

## Abstract

Since their discovery, extracellular vesicles (EVs) have changed our view on how organisms interact with their extracellular world. EVs are able to traffic a diverse array of molecules across different species and even domains, facilitating numerous functions. In this study, we investigate EV production in *Haloferax volcanii*, as representative for Euryarchaeida. We uncover that EVs enclose RNA, with specific transcripts preferentially enriched, including those with regulatory potential, and conclude that EVs can act as an RNA communication system between haloarchaea. We demonstrate the key role of an EV-associated Ras superfamily GTPase for EV formation in *H. volcanii* that is also present across other diverse evolutionary branches of Archaea. Ras superfamily GTPases are key players in eukaryotic intracellular vesicle formation and trafficking mechanisms that have been crucial for the emergence of Eukaryotes. Therefore, we propose that archaeal EV formation could reveal insights into the origin of the compartmentalized eukaryotic cell.

## INTRODUCTION

Extracellular vesicles (EVs) (also referred to as outer membrane vesicles, membrane vesicles, or exosomes) are small membrane bound structures that bud off from the cellular envelope, and are produced by living cells across all domains of life [1–3]. They are able to enclose a wide range of cargo, including proteins, nucleic acids, and signaling molecules, facilitating a mechanism of interaction with the extracellular world. Communication mediated through EVs provides specific advantages for the cell, such as protection of the cargo from environmental stressors and degradation, the concentration of specific molecules into a self-contained structure, and the potential for selective delivery to designated targets [4, 5]. With the diversity of EV composition and the advantages of EV-based communication, prokaryotic EV trafficking has been connected to a wide range of cellular functions. EVs have been discovered to act as defense against viral infection and antibiotic stress [6], mediating Bacteria-host interactions through the trafficking of regulatory RNA [7, 8], and facilitating the transfer of genetic material between cells [9, 10]. Both their ubiquity amongst organisms and cellular functions make EVs an exciting new field for exploring intercellular communication and expand our view of the dynamics driving microbial environments.

EVs are known to be present in marine and aquatic samples [11, 12], and they likely play important roles in regulating environmental microbial populations. However, while there is a growing amount of research focusing on EVs deriving from pathogenic Bacteria and their role in Bacteria-host interactions, fewer studies have investigated the role that EVs play in microbial ecology, and even less investigate EVs in Archaea. Within the archaeal domain, EVs from only a few organisms have been studied. For the *Thermoproteota* (formerly *Crenarchaeota*) genus, *Sulfolobus*, vesicles were found to enclose proteinaceous toxins [13] as well as fragmented genomic DNA [14]. Members of the Euryarchaeida have also been found to produce EVs enriched with DNA such as *Thermococcus* [15]*. Halorubrum lacusprofundi* [16] was found to produce specialized EVs including plasmid-encoded proteins and plasmid DNA (named plasmid vesicles, PVs). These studies demonstrate the ability of archaeal EVs to transport DNA between cells, which suggests that EV production may play an important role in horizontal gene transfer in Archaea. This route of genetic exchange may be especially beneficial for organisms in extreme environments to counteract various common stressors.

EV production in *Sulfolobus* has been linked to its ESCRT (endosomal sorting complex required for transport)-like cell division machinery [14]. The ESCRT system is well studied in Eukaryotes, and is responsible for the sorting and production of exosomes and the budding of various viruses [17], suggesting an evolutionary linkage between EV formation in *Thermoproteota* and exosome production in Eukaryotes. However, ESCRT-like proteins are not present in most currently annotated Euryarchaeida genomes [18], suggesting that a different mechanism is responsible for EV production. Proteins homologous to eukaryotic intracellular vesicle trafficking system were identified in PVs from *Hrr. lacusprofundi*, a member of the Euryarchaeida, implying that multiple mechanisms of vesicle production exist among the archaeal domain [16].

In order to understand EV production in Euryarchaeida, and in particular halophilic Archaea (haloarchaea), we used the model organism, *Haloferax volcanii*, to investigate the composition of EVs as well as their capacity to transfer their cargo to other organisms. We observe for the first time particular RNAs being enriched in EVs of various haloarchaea, and demonstrate that the RNA cargo can be transferred between cells of the same species. We also investigated the roles of various genes in EV production, suggesting a mechanism for EV generation in halophilic Archaea that is related to intracellular vesicle trafficking in Eukaryotes. From our findings, we hypothesize that halophilic Archaea utilize EVs to communicate and potentially regulate the microbial community in hypersaline environments.

## RESULTS

### EV production in H. volcanii is dependent on growth conditions

The production of EVs and PVs (specialized EVs in the presence of the plasmid, pR1SE) has been reported for the Haloarchaeon, *Halorubrum lacusprofundi* [16]. Both were characterized, revealing the packaging of the plasmid pR1SE and a number of plasmid proteins for PVs, and a set of EV-associated proteins for EVs in absence of pR1SE. To further investigate the generation and potential function of EVs in Haloarchaea, we chose to use *H. volcanii* because it is a well-established model organism for haloarchaeal cell biology with a number of genetic tools available [19, 20]. The capability of *H. volcanii* to produce EVs was also previously reported under UV irradiation [21].

EVs were isolated from culture supernatants of *H. volcanii* and were observed to be circular with a diameter ranging from 50 to 150 nm (Figure 1). Purification of EVs by iodixanol (OptiPrep™) density-based gradient purification (see Methods) resulted in EVs concentrating into two distinct bands in the gradient (Supplementary Figure 1A and B). No obvious differences distinguishing the two bands could be observed by TEM (Supplementary Figure 1C and D).

**Figure 1.**
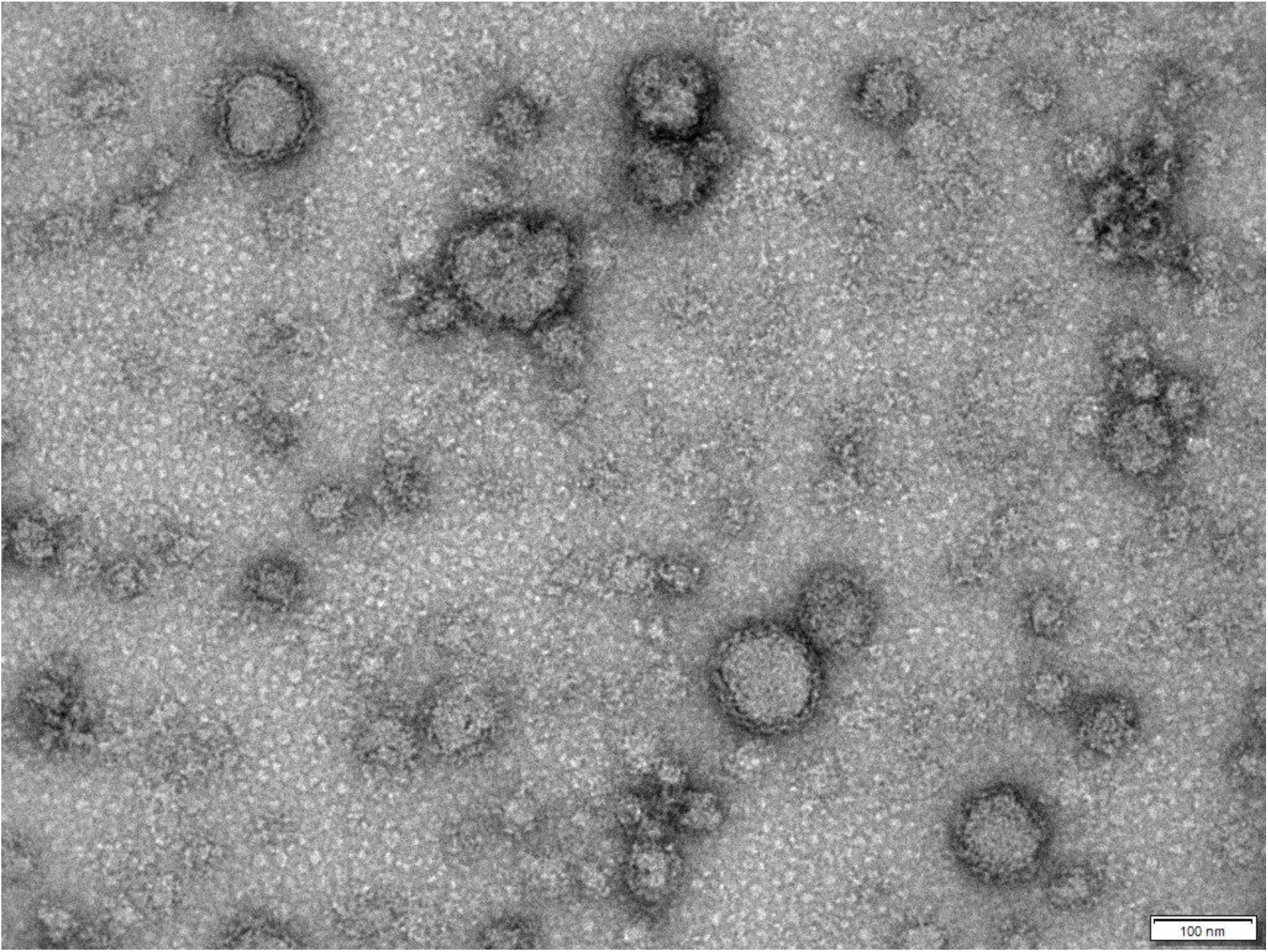
Transmission electron micrograph of EVs from *H. volcanii DS2.* EVs were isolated from culture supernatants and purified by density gradient centrifugation. Size bar represents 100 nm.

Initial efforts to isolate EVs close to the documented optimal temperature of *H. volcanii* at 45 °C yielded low amounts of EVs. However, when lowering the temperature of growth to 28 °C, we were able to isolate EVs from culture supernatants, suggesting that EV production is temperature dependent. To determine the optimal conditions for EV production, we tested different growth temperatures using a fluorescence-based method for EV quantification (see methods), and confirmed a temperature-dependent production of EVs reflecting what we observed in large scale purifications (Figure 2A). The abundance of EVs in the supernatant was determined to peak during early stationary phase (Figure 2C, Supplementary Figure 2C), indicating that the rate of EV production versus uptake varies between growth stages. Since stress has also been reported to induce EV production, we tested environmental stress conditions such as UV exposure and virus infection using immunodetection-based EV quantification (see methods). Both UV stress (p-value = 0.067) and infection with the chronic infecting virus, HFPV-1 [22] (p-value = 0.130), did not appear to alter EV production significantly under the conditions tested (Figure 2B).

**Figure 2.**
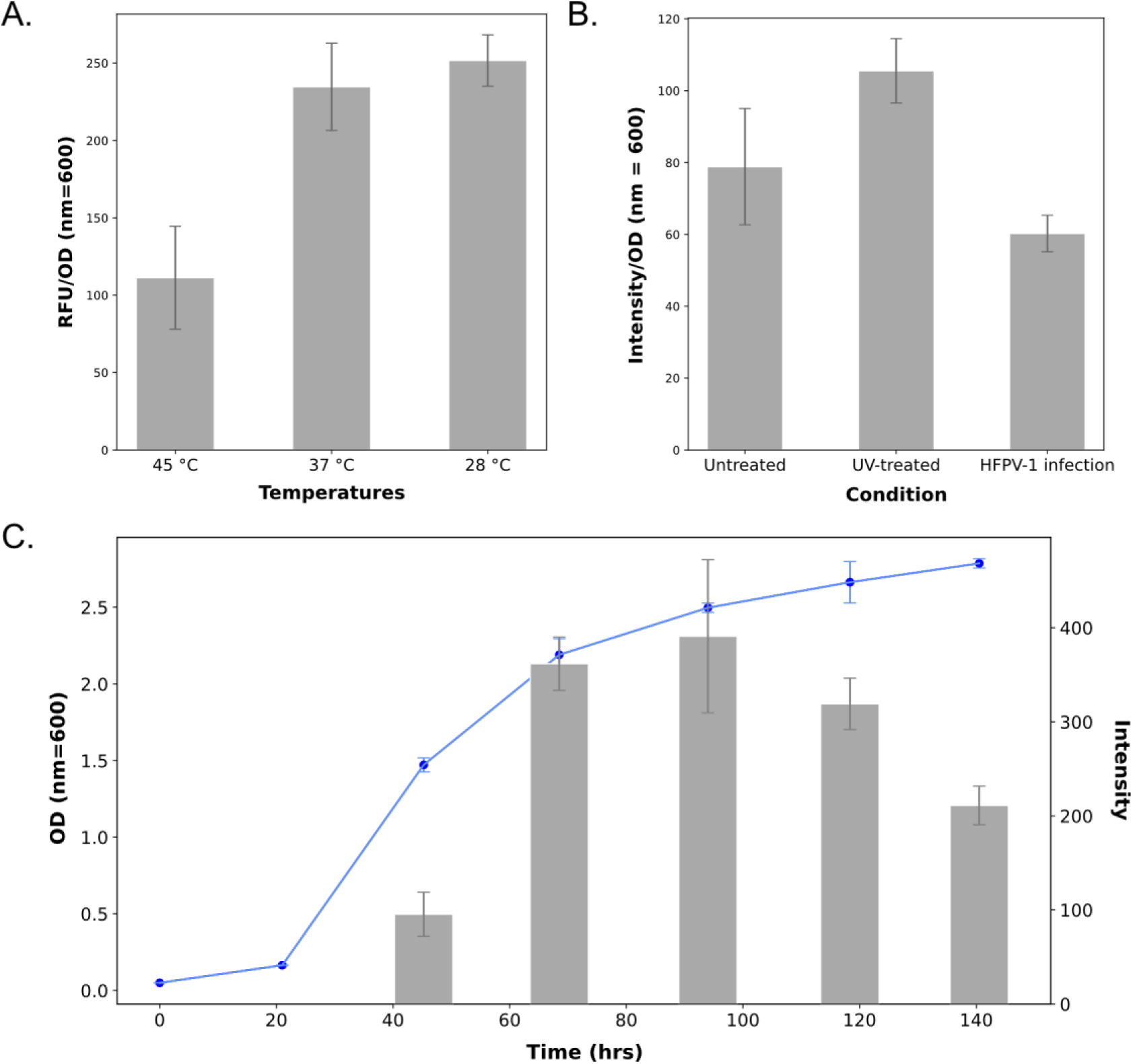
EV abundancies under different conditions and stages of growth. EVs were quantified (gray bars) from culture supernatants of *H. volcanii DS2* grown under different temperatures **(A)**, exposure to UV radiation and viral infection **(B)**, and during different stages of growth **(C)**. Blue line illustrates the OD (nm = 600) of cell cultures averaged over 3 replicates. EV quantities from **(A)** were quantified using fluorescence (see methods) and **(B)** and **(C)** were quantified using immunodetection (see methods). EV quantifications at different temperatures and different conditions **(A and B)** were conducted at stationary phase, and the EV signal was normalized to OD (nm = 600) at the point of measurement. Original spot blots are presented in Supplementary Figure 2. Error bars represent standard deviation over three replicates. RFU = relative fluorescence unit.

### H. volcanii EVs are associated with RNA

EVs of both *Sulfolobus* (*Thermoproteota*) and *Thermococcus* (Euryarchaeida) were previously shown to enclose DNA [14, 15]. To determine the nucleic acid contents of *H. volcanii* EVs, we attempted to isolate both DNA and RNA from a purified EV preparation. While DNA extraction yielded negligible amounts of DNA associated with the EVs, RNA extraction revealed high yields of EV-associated RNA. Nuclease (DNase and RNase) treatment of the EVs prior to RNA extraction did not eliminate the presence of RNA, confirming that the transcripts are protected and likely enclosed within the vesicles. Analysis of the size distribution of the enclosed RNA revealed differences between EV-associated RNA and intracellular RNA (Figure 3A). While ribosomal subunits were prominent in both EV and cellular preparations, we observed populations of RNAs that are significantly enriched in EVs with a tendency towards smaller transcripts (Supplementary Figure 3).

**Figure 3.**
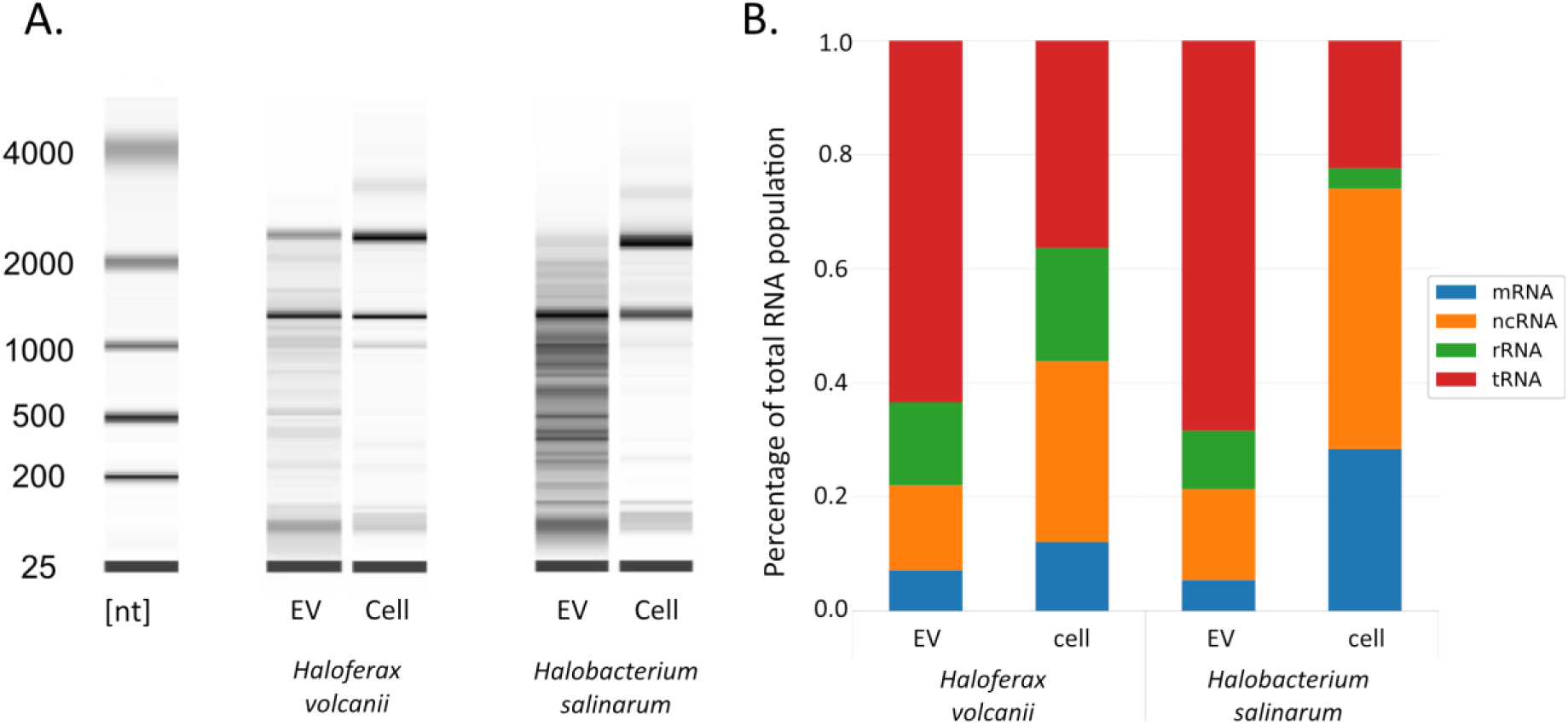
RNA composition of haloarchaeal EVs. **(A)** Analysis of the size distribution of RNA extracted from purified EVs and whole cells of *H. volcanii* and *Hbt. salinarum*. **(B)** Distributions of RNA subpopulations comparing cellular and EV-associated RNAs for *H. volcanii* and *Hbt. salinarum*.

### EV-associated RNA is enriched in tRNAs, rRNAs and ncRNAs

Preliminary sequencing approaches of EV-associated RNA (see Supplementary Results) revealed that using small RNA libraries best reflects the RNA content of EVs. Additionally, we compared the RNA content of upper and lower EV bands in density gradients, revealing that the RNA content alone is unlikely the major factor leading to two subpopulations of EVs. Finally, we deciphered that comparing EV-associated and intracellular RNA is crucial to determining the nature of RNA preferentially associated with EVs.

Using small RNA libraries, we identified around 4,400 genes represented by EV-associated transcripts, comprising the majority of the *H. volcanii* genome with around 79.5% ± 10.5% of genome covered by at least one read (85.2% ± 0.8% for intracellular reads). Though this encompasses nearly all genes in the *H. volcanii* genome, only 474 of the transcripts identified had a TPM (transcript per million) greater than 10, suggesting the majority of identified EV-associated RNA can be considered transcriptional noise. The most abundant of the identified transcripts were tRNAs (68.9 ± 2.1%), followed by non-coding RNAs (ncRNA, transcripts that do not encode a protein, excluding rRNA and tRNA) and rRNAs (16.1 ± 0.9% and 10.4 ± 1.2% respectively) (Figure 3B). The identified ncRNA include intergenic sRNAs [23, 24] and antisense RNAs (asRNA). While we also detected mRNAs in the EV fraction, they only constitute about 4.6 ± 0.1% of the RNA population.

Notably, when we normalized the EV-associated RNA to the intracellular RNA (see Methods), the EV-associated RNA represented a unique subset of transcripts with little variation among replicates (Figure 4A, Supplementary Table 6). We identified 230 transcripts as highly abundant (TPM > 10) and highly enriched (log2 > 1) in EVs. This population comprised of tRNAs, rRNAs, ncRNAs and mRNAs, with tRNAs being the most dominant group. Surprisingly, while the mRNA fraction was the least represented among EV-associated RNA, the most enriched (242-fold) among all transcripts was the mRNA for the S-layer protein (HVO_2072). A Northern blot analysis probing for the full-length mRNA (gene length 2484 bp) in intracellular and EV-associated RNA revealed only smaller fragments of the transcript associated with EVs (Supplementary Figure 4). Besides HVO_2072, the remainder of highly enriched mRNAs were low in abundance (TPM < 10). We predict that the enrichment of these mRNA relates to their proximity to the cell envelope (transcripts of membrane associated transporters and other transmembrane proteins). Both rRNA and tRNA were previously reported to be associated with EVs from bacterial and eukaryotic organisms [8, 25, 26], and we hypothesize that their presence across RNA-containing EVs is likely due to their structural stability and abundance within cells.

**Figure 4.**
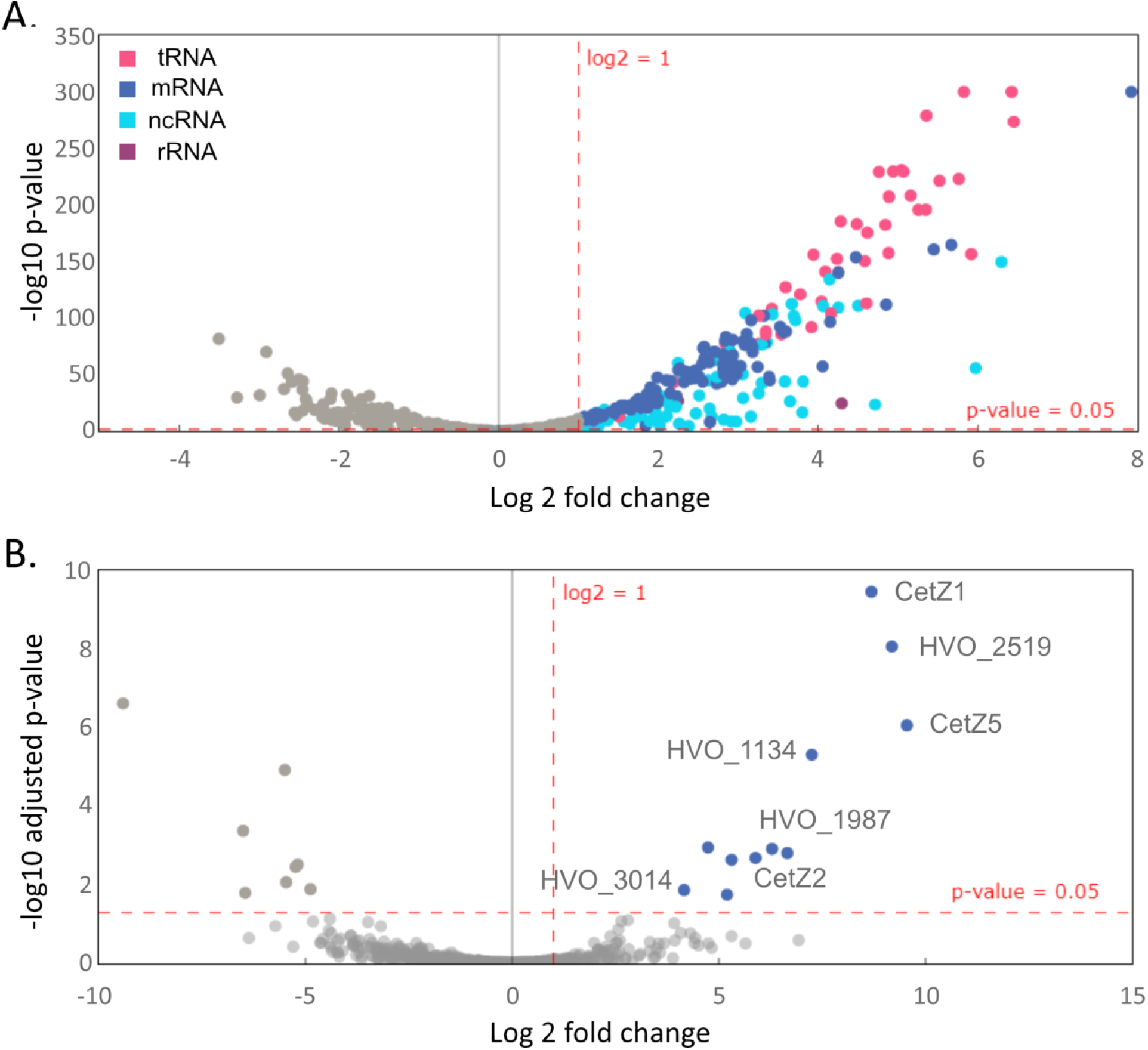
EV associated RNA and proteins. Volcano plots of RNA (**A**) and protein (**B**) abundances in EVs after normalization to cellular RNA abundances and protein abundances from cell membranes. Differential RNA abundancies and p-value calculated using DESeq2. Differential protein abundancies and adjusted p-values calculated by DEP (see methods). Raw data are presented in Supplementary Tables 6 and 10.

Within the population of ncRNAs associated with EVs, we identified 74 ncRNAs that are both highly abundant and enriched in EVs (Table 1, Supplementary Table 7). This population consists of intergenic RNAs as well as asRNAs. However, no function has been predicted for any of the intergenic ncRNAs so far. We also screened the ncRNAs for consensus sequences or a common secondary structure as specific selection markers for EV packaging; however, no common motif could be identified. Nevertheless, the identified asRNAs (21 asRNAs) appeared to exhibit sequence and structural similarities (Supplementary Figure 5A and B). The average length of these asRNAs was 45.5 nt (± 5.8 nt), and all are associated with the 5’ end of ISH3-, ISH5-, ISH8-, ISH9- and ISH11-type transposases from across the genome, overlapping with the predicted start codons of the respective transposase (Supplementary Figure 5C).

**Table 1.**
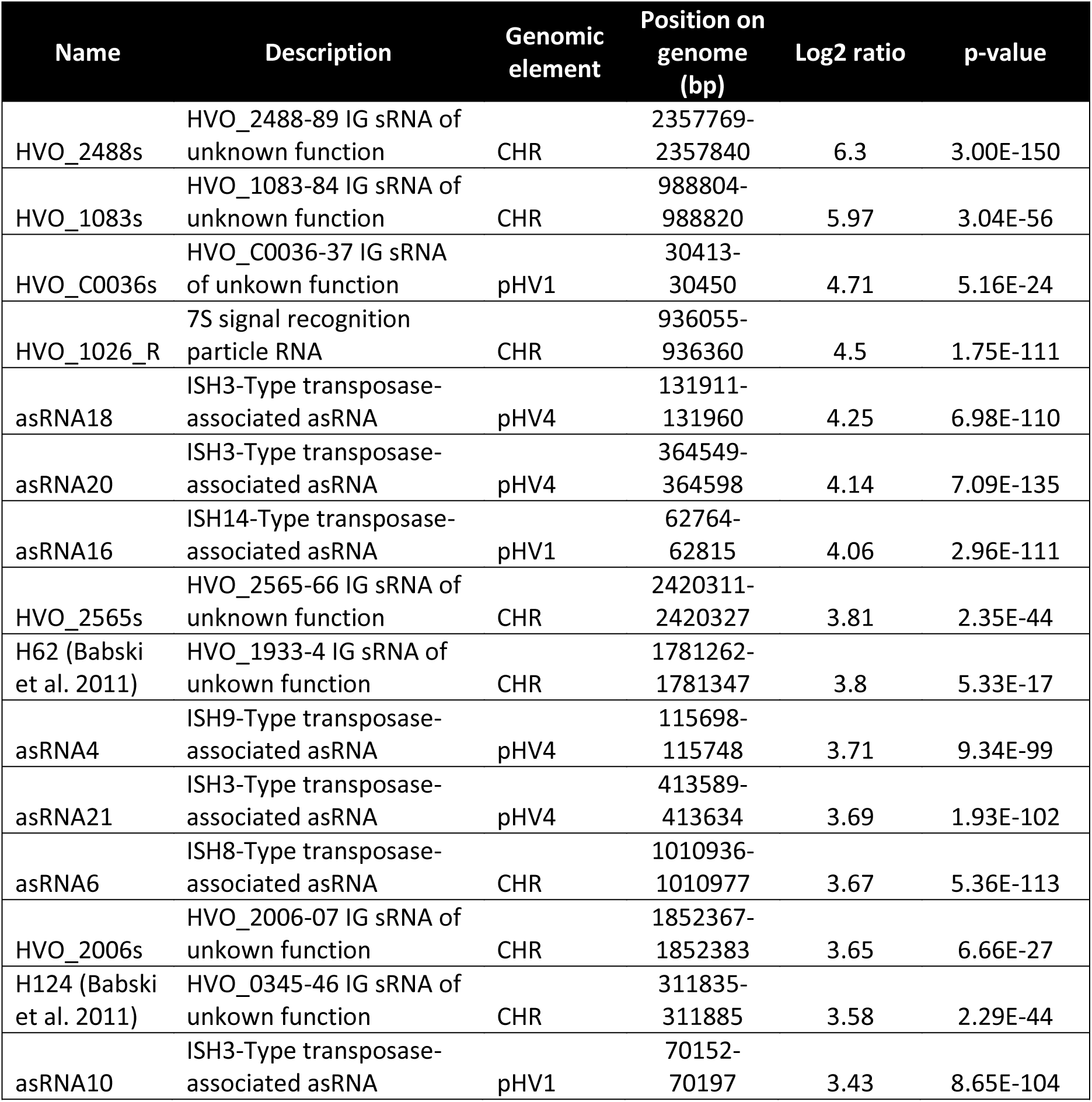
Top 15 noncoding RNAs enriched in EVs of *H. volcanii*. Full transcriptomic data set in Supplementary Table 6. (IG = intergenic, CHR = Chromosome, pHV1 – pHV4 = plasmids)

| Name | Description | Genomic element | Position on genome (bp) | Log2 ratio | p-value |
| --- | --- | --- | --- | --- | --- |
| HVO_2488s | HVO_2488-89 IG sRNA of unknown function | CHR | 2357769-2357840 | 6.3 | 3.00E-150 |
| HVO_1083s | HVO_1083-84 IG sRNA of unknown function | CHR | 988804-988820 | 5.97 | 3.04E-56 |
| HVO_C0036s | HVO_C0036-37 IG sRNA of unknown function | pHV1 | 30413-30450 | 4.71 | 5.16E-24 |
| HVO_1026_R | 7S signal recognition particle RNA | CHR | 936055-936360 | 4.5 | 1.75E-111 |
| asRNA18 | ISH3-Type transposase-associated asRNA | pHV4 | 131911-131960 | 4.25 | 6.98E-110 |
| asRNA20 | ISH3-Type transposase-associated asRNA | pHV4 | 364549-364598 | 4.14 | 7.09E-135 |
| asRNA16 | ISH14-Type transposase-associated asRNA | pHV1 | 62764-62815 | 4.06 | 2.96E-111 |
| HVO_2565s | HVO_2565-66 IG sRNA of unknown function | CHR | 2420311-2420327 | 3.81 | 2.35E-44 |
| H62 (Babski et al. 2011) | HVO_1933-4 IG sRNA of unknown function | CHR | 1781262-1781347 | 3.8 | 5.33E-17 |
| asRNA4 | ISH9-Type transposase-associated asRNA | pHV4 | 115698-115748 | 3.71 | 9.34E-99 |
| asRNA21 | ISH3-Type transposase-associated asRNA | pHV4 | 413589-413634 | 3.69 | 1.93E-102 |
| asRNA6 | ISH8-Type transposase-associated asRNA | CHR | 1010936-1010977 | 3.67 | 5.36E-113 |
| HVO_2006s | HVO_2006-07 IG sRNA of unknown function | CHR | 1852367-1852383 | 3.65 | 6.66E-27 |
| H124 (Babski et al. 2011) | HVO_0345-46 IG sRNA of unknown function | CHR | 311835-311885 | 3.58 | 2.29E-44 |
| asRNA10 | ISH3-Type transposase-associated asRNA | pHV1 | 70152-70197 | 3.43 | 8.65E-104 |

### Analysis of EV-associated RNA under infection with a virus reveals viral transcripts associated with EVs

Direct interactions between EVs and viruses have been documented, demonstrating the capacity for EVs to act as a viral defense mechanism [6] or facilitate viral propagation [27]. While we did not observe significant changes to EV quantities under infection with HFPV-1 (Figure 2B), we wanted to test whether infection with HFPV-1 would influence the RNA composition of EVs and thereby possibly indirectly influence virus-host interactions.

While it was shown that infection with HFPV-1 drastically altered the transcriptomic landscape of the cell during exponential and early stationary growth [22], sRNAseq in late exponential growth revealed a nearly identical transcriptional profile when comparing infected and uninfected cells (Supplementary Fig. 6A). Only two genes showed a significant upregulation (log2 > 1) in the infected cells, HVO_2657 and HVO_0272; however, both are in general rather weakly expressed (TPM < 15). When comparing the RNA content of EVs between infected and uninfected cells, two transcripts were found to be significantly higher in abundance in EVs of infected cells: HVO_A0466 and HVO_0272 (Supplementary Fig. 6A). While HVO_0272 mRNA was about 4-fold upregulated in infected cells (log2 ∼ 2), it was about 10-fold upregulated (log2 ∼ 5) in EVs of infected cells (Supplementary Fig. 6), indicating that the packaging of this transcript into EVs increases significantly upon infection. Surprisingly, it appeared that the majority of reads mapping to HVO_0272 only map to two short regions of about 30 nt within the coding region of the gene that are identical to a region on the viral genome. Therefore, we conclude that the upregulation of HVO_0272 is due to viral transcripts mapping to the host genome.

Subsequently, when mapping reads to the virus genome, we detected a significant amount of viral transcripts in EVs. While only 1.7 ± 0.07% of intracellular RNA mapped to the HFPV-1 genome, 4.0 ± 0.10% of EV-associated RNA mapped to the viral genome, suggesting a slight enrichment of virus-derived transcripts in EVs. Both cellular and EV-associated RNA mapped the entire HFPV-1 genome, and no enrichment of particular viral RNAs could be detected in EVs (Supplementary Figure 7). However, the detection of viral transcripts within EVs shows that they are also exported in EVs together with host RNA.

### Generation of RNA-enriched EVs could be detected in other haloarchaea

We were also interested in whether RNA-enriched EVs are a conserved phenomenon among halophilic Archaea. Therefore, we tested *Halobacterium salinarum* and *Halorubrum lacusprofundi* for EV production and EV-associated RNA content. EVs could be isolated from both organisms (Supplementary Figure 8A and B), and they were likewise found to be enriched in RNA. The size distribution of EV-associated RNA indicates an enrichment for a specific RNA population (Figure 3A and Supplementary Figure 8C). However, we only proceeded with sequence analysis of EV-associated RNA of *Hbt. salinarum*.

RNA sequencing revealed 85.4% of the *Hbt. salinarum* genome was covered by at least one read from EV-associated RNA (94.5% from intracellular RNA library). The distribution of RNA populations were very similar between *H. volcanii* and *Hbt. salinarum* EVs (Figure 3B), with the majority of EV-associated transcripts being tRNAs.

After normalizing EV-associated RNA with intracellular RNA, we identified 228 transcripts as highly abundant and highly enriched in *Hbt. salinarum* EVs (Supplementary Table 8). The transcript for the S-layer subunit was similarly one of the most enriched EV-associated transcripts in *Hbt. salinarum*. The most enriched transcript was a 29 nt asRNA mapping to the coding region of VNG_RS00640, a predicted helix-turn-helix domain protein of unknown function. We also identified 16 highly enriched transposase-associated asRNA that associate with a larger range of transposase families than those from *H. volcanii*, some of which overlap with the respective predicted start codon. In total, 35 ncRNAs were identified as highly enriched and highly abundant in EVs of *Hbt. salinarum*. Of the ncRNA enriched in *Hbt. salinarum* EVs, 6 are sense-overlapping transposase-associated RNA (sotRNA) [28], and 2 are intergenic sRNAs with high sequence identities to the predicted sRNAs from *H. volcanii*, HVO_2908s and H3.2 [23], that were also found in *H. volcanii* EVs.

### EVs are enriched with specific proteins

The protein compositions of *H. volcanii* EVs and their respective cellular membranes were identified by mass spectrometry (LC-MS). Since we determined no difference between the RNA content of upper and lower bands of EV preparations in OptiPrep™ gradients, we analyzed the bands separately for protein content to uncover possible differences in protein composition. We identified 401 proteins associated with EVs deriving from the upper band and 384 proteins from the lower band. However, only 1 protein was found to be more abundant in the upper band and 2 more abundant in the lower band, while the majority of proteins appeared to be consistent between upper and lower bands (Supplementary Figure 9). Therefore, we concluded that protein content alone is most likely not the major factor causing the separation into two bands and pooled the results from both bands for further analysis.

In total, we identified 328 proteins associated with EVs and 668 proteins in the cellular membrane preparations. We compared the abundancies of proteins in EVs with those in cell membranes and obtained 11 proteins significantly enriched in EVs (log2 > 1, adjusted p-value < 0.05) (Supplementary Table 9, Figure 4B), including one protein exclusively detected in EVs (hypothetical protein, HVO_2519, with unknown function and no detectable conserved domains). Several CetZ proteins, including CetZ5 (HVO_2013), CetZ1 (HVO_2204), and CetZ2 (HVO_0745), were also identified to be enriched in EVs. CetZ1 and CetZ2 have been shown to be involved in controlling cell shape and motility in *H. volcanii*, and the CetZ protein family has been predicted to be involved in other cell surface-related functions in Archaea [29, 30].

Other highly enriched, notable proteins include FtsZ2 (cell division protein), HVO_1134 (hypothetical protein), HVO_2985 (hypothetical protein), HVO_1964 (YlmC sporulation protein), and two ABC transporter proteins. Hypothetical protein, HVO_1134, is predicted to contain a rod domain (Interpro) [31], building α-helical coiled coils (Phyre2, I-TASSER) [32, 33], a structure that is also found in viral fusion proteins, representing a candidate gene potentially involved in EV fusion.

Most interesting was the enrichment of the small GTPase, HVO_3014 (OapA), a homolog of the Sar1/Arf1-like GTPase that was also found to be enriched in *Hrr. lacusprofundi* EVs, Hlac_2746 [16] (Figure 4B). OapA was initially thought to have an influence on genome replication due to its association with the origin of replication. However, despite a study characterizing a mutant strain, no distinct function could be assigned to OapA so far [34]. The best homologs for HVO_3014, using a Hidden Markov Model (HMM) based search [35] are Rab-family GTPases (Supplementary Figure 10). Rab-family GTPases belong to the Ras superfamily of small GTPases whose functions include regulating vesicle and membrane trafficking in Eukaryotes[36]. In Eukaryotes, all three types of carrier vesicles (COPI, COP II and clathrin-coated vesicles) require the activation of a small Ras family GTPase to initiate vesicle coat assembly [37]. A COPI-like mechanism of vesicle formation involving a Ras family GTPase was already proposed for *Hrr. lacusprofundi* [16].

We also analyzed the protein composition of EVs from UV-treated cells and compared them with membrane-associated proteins isolated from their respective cells, to determine whether UV treatment would alter protein composition of the EVs. We identified 377 proteins associated with EVs and 668 proteins associated with their respective cell membranes. We identified 11 proteins to be enriched in EVs from UV-treated cells (Supplementary Table 10, Supplementary Figure 11A). All proteins identified as enriched in EVs from untreated cells were also identified as enriched in EVs from UV-treated cells, except for the small GTPase, HVO_3014, that was calculated as equally enriched but did not pass the E-value threshold. Instead, one additional ABC transport protein (HVO_2399) was identified as enriched.

Differential expression analysis only identifies proteins that are present in higher abundancies in EVs than in cell membranes, leaving out other proteins that could be functionally relevant but are present in equal or lower abundancies when normalized to the cell membrane. For instance, the small GTPase, HVO_3014, was not identified to be enriched in EVs from UV-treated cells using a standard threshold, yet we observe its integral relationship to EV production in *H. volcanii* (see below). Therefore, we also identified the proteins that were found to be present amongst all 12 EV samples analyzed (Supplementary Table 11) and identified 285 proteins present across all samples. All proteins identified as enriched by differential expression analysis were also present in this list, except HVO_2399 identified as enriched in EVs from UV-treated cells, suggesting that the protein composition slightly changes upon UV exposure. The most abundant protein was cytoskeletal protein, CetZ1, followed by the S-layer protein, HVO_2072. Other notable proteins within this list were ribonuclease J (HVO_2724), diadenylate cyclase (HVO_1660), and HVO_1020 (hypothetical protein). RNase J is an exonuclease, and could be relevant to the enrichment of RNAs found associated in the EVs. Diadenylate cyclases are responsible for the production of cyclic-di-AMP, a common secondary messenger among Bacteria and Archaea, including *H. volcanii* [38]. HVO_1020 is a homolog (55% sequence identity) to *H. lacusprofundi* PQQ ß-propeller repeat domain-containing protein, Hlac_2402, that was also identified in *H. lacusprofundi* EVs [16] and has similarities to the gamma subunit of COPI.

### Knockout of oapA abolishes EV formation

To investigate the proposed involvement of OapA in EV production in *H. volcanii*, we compared the phenotypes of an OapA knockout strain [34] to the respective parental strain (H26). The OapA knockout strain yielded a dramatically reduced amount of EVs in comparison to the parental strain (about 70% reduction, p-value = 0.007) (Figure 5A, Supplementary Figure 2F).

**Figure 5.**
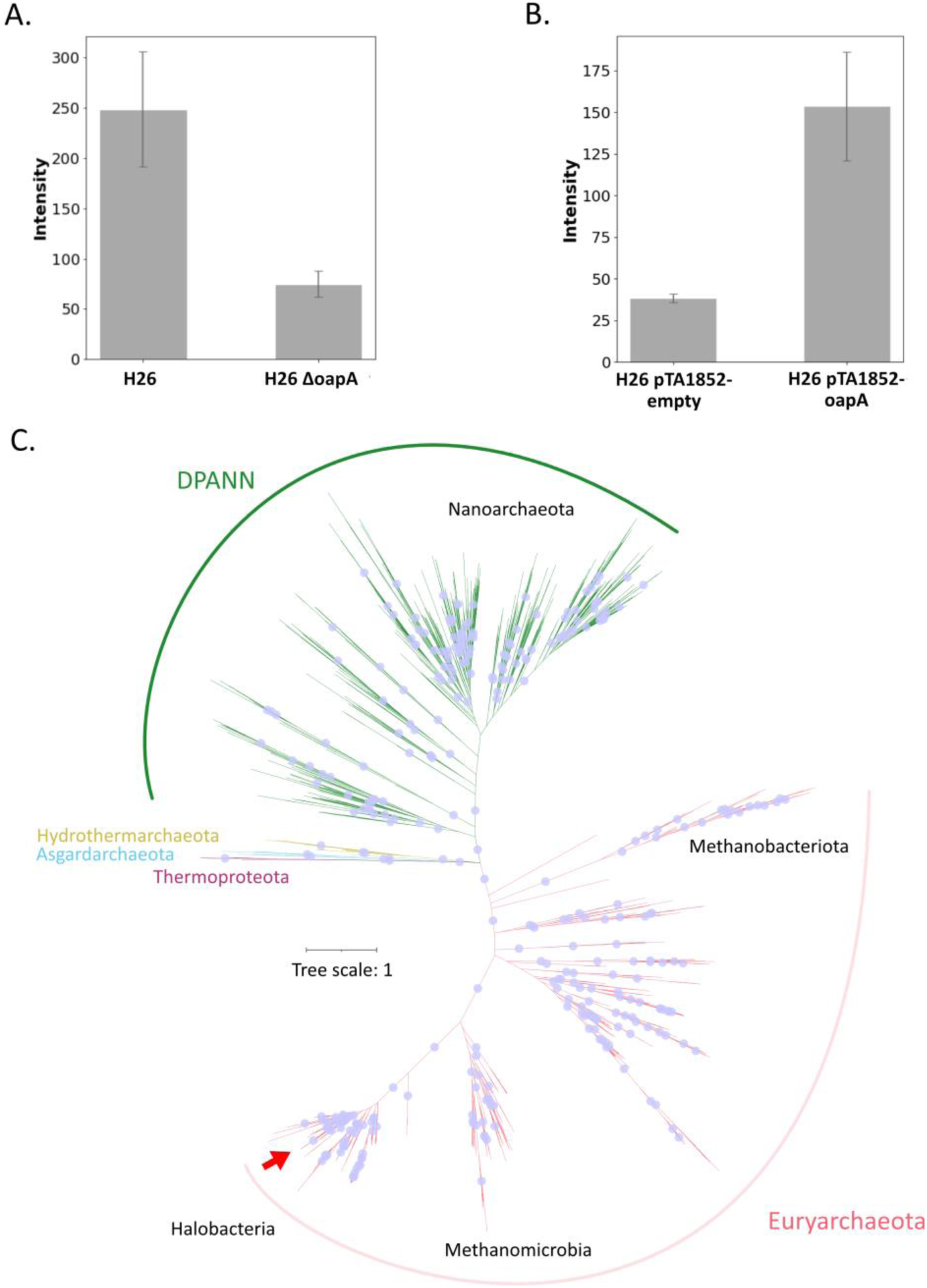
Analysis of *H. volcanii* EV-associated GTPase, OapA, and homologues in other Archaea. **(A)** Quantification of EVs in the culture supernatant of the OapA knockout strain and the respective parental strain. **(B)** Quantification of EVs in the culture supernatant of OapA knockout strain and respective parental strain expressing OapA_t_ under an inducible promotor (pTA1852-OapA_t_), compared to a control with the empty vector (pTA1852-empty). EVs were quantified using immunodetection (see methods) and were averaged over three replicates with error bars denoting standard deviations. Original spot blots are presented in Supplementary Figures 2D and 16A. **(C)** Unrooted phylogenetic tree of the identified small GTPases across the archaeal domain. Red arrow indicates position of GTPase from *H. volcanii*. Blue dots represent branches with bootstrap value greater than 95.

Gradient purification of concentrated OapA mutant supernatant resulted in either no distinct band or only one band with reduced intensity at about the same height of the upper band in density gradients of the parental strain (Supplementary Figure 12). RNA extracted from this single band yielded very low RNA concentrations and was not detectable on a fragment analyzer. Interestingly, resequencing of the strain revealed the activity of a known proviral region (HVO_1422 – HVO_1434) in the *H. volcanii* genome [39] (Supplementary Figure 13). Therefore, we propose that the remaining particles isolated from supernatants of OapA knockout strain cultures that concentrated in one band of the density gradient could be cell debris and virus particles, and that the strain is unable to produce EVs associated with RNA. Phenotypic changes of the cell shape/morphology were also observed for the knockout strain (Supplementary Fig. 14A). The formation of rod-shaped cells appears to be less frequent when *oapA* is deleted. Interestingly, the OapA mutant also showed a slightly increased growth rate when compared to the parental strain in our laboratory (Supplementary Fig. 14B).

Further, overexpression of OapA in an OapA wild-type background strain (H26) resulted in increased vesicle production (4-fold, p-value = 0.005) (Figure 5). The hypervesiculation phenotype was confirmed by both quantification and TEM (Figure 5, Supplementary Figure 15B and C), further implicating the crucial role of OapA for EV production.

### Ras-superfamily GTPases are conserved amongst various archaeal clades

To get an overview of whether this particular Ras-superfamily GTPase is present in other Archaea, we searched for proteins with high similarity to HVO_3014 against archaeal and bacterial GTDB species representatives using an alignment score ratio approach (see methods) [40]. 1666 archaeal proteins were identified across 14 phyla of Archaea, with an uneven distribution of GTPases across these phyla (Supplementary Figure 16, Supplementary Table 12). The majority of GTPases were identified among the Euryarchaeida, *Halobacteria* and *Methanobacteriota*, as well as in 7 DPANN phyla, including *Nanoarchaeota*, *Nanohaloarchaeota*, and *Altiarchaeota*. Interestingly, only 8 *Korarchaeota* out of 970 *Thermoproteota* genomes analyzed contained a homologous small GTPase. Further, only 8 out of 183 *Asgardarchaeota* were identified to contain a homologous small GTPase. As expected, we were also unable to identify any homologs in two well-studied EV-producing organisms, *Sulfolobus* (*Thermoproteota*), that is known to generate EVs using ESCRT-like proteins, and *Thermococcus* (*Methanobacteriota_B*), for which other proteins were suggested to be involved in EV formation [14, 41].

A phylogenetic tree was constructed from the alignment of the GTPase database to determine the evolutionary relationships between the GTPases identified and whether they align with taxonomy of the organisms (Figure 5C). Both DPANN and Euryarchaeota appear to have distinct subclades of this family of GTPase that group separately from each other. Within each superphyla, there was also some internal organization. For instance, all of the GTPases of *Halobacteria* and *Methanobacteriota* formed distinct groups, separate from the other Euryarcheaeida with mostly small branches, suggesting strong sequence similarity within these groups. Similarly, the nanoarchaeal GTPases also formed a distinct group from the other DPANN organisms. However, the branch lengths were long in relation to the rest of the tree, suggesting a higher variation of sequences within this group. While the bootstrap values of the DPANN and Euryarchaeida branches (100 and 99 respectively) indicate that this division in this family of GTPases based on taxonomy is well supported, the organization of the smaller third branch containing *Asgardarchaeota*, *Thermoproteota*, and *Hydrothermarchaeota* were not as well supported (bootstrap value of 56). However, the GTPases within this cluster form smaller sub-clusters that correlate with their phyla.

### Testing other knockout mutants provides further insight into the mechanisms of EV formation

CetZ1 and CetZ2 were amongst the most abundant proteins in EVs; however, we were able to isolate EVs from the supernatant of both CetZ1 and CetZ2 knockout strains. While quantification of EVs by the immunodetection-based method was not possible for the CetZ1 knockout strain, we did not have any indication when purifying EVs that EV production was drastically altered for CetZ1 and CetZ2 knockout strains (Supplementary Figure 17). RNA could also be isolated from EVs of this strain, and the size distribution of EV-associated RNA was nearly identical when compared to the parental strain (Supplemental Figure 17).

Previous studies in Bacteria have shown that destabilization of the cell envelope results in a ‘hypervesiculation’ phenotype [42, 43]. To investigate whether changes in cell envelope stability would similarly affect EV production in *H. volcanii*, we assessed EV production in an *aglB* knockout strain. Cells lacking AglB are unable to N-glycosylate the S-layer glycoprotein and absence of AglB results in enhanced release of the S-layer glycoprotein [44]. Thus, deletion of this protein causes a destabilization of the structural integrity of the cell envelope. Indeed, we observed a significant increase in EV production from the *aglB* knockout strain during the purification process as well as by TEM (Supplementary Figure 18A and B). While we could not confirm this result when using the immunodetection-based assay for quantifying EVs (Supplementary Figure 18D), EV quantification by fluorescence staining confirmed an increase in EV production (Supplementary Figure 18E), indicating that CetZ1 incorporation into EVs is altered in this mutant. Interestingly, we observed a drastic change to the morphology of EVs in the mutant. The surface of EVs isolated from the *aglB* knockout strain was significantly different from EVs of the parental strain, appearing very fuzzy (Supplementary Figure 18C), likely due to the instability of the S-layer. Further, while we isolated a significantly larger amount of EVs from the mutant, the RNA yield was slightly lower in EVs (Supplementary Figure 18F), suggesting that a great portion of EVs are likely budding randomly without enclosing RNA.

### Lipid analysis reveals differences in the relative abundance of distinct lipids between cells and EVs

To determine whether EVs selectively enclose particular lipids, we analysed the lipid content of EVs and compared the relative abundances of different lipid compounds in EVs to that of cell membranes and total cells of *H. volcanii*. We detected minimal differences in proportions of lipid types between EVs deriving from upper and lower bands of a density gradient (Supplementary Figure 19B), indicating that the lipid content alone is not the differentiating factor between the two subpopulations. We therefore chose to pool samples from different bands of each replicate for comparison.

Lipids with phosphate-based polar head groups were dominant across all samples. Methylated-phosphatidylglycerolphosphate-archaeol (Me-PGP-ARP) both in saturated and unsaturated form (:n) represented the most abundant lipid across samples, with relative abundances of 53.9 ± 2% in whole cells, 66.4 ± 8.81% in cell membranes and 46.8± 1.91% in EVs (Figure 6, Supplementary Figure 19A). The ratio of unsaturated to total Me-PGP-AR abundance was identical in cells and cell membranes (28 ± 1.64% and 29 ± 3.6%), but the comparative amount of unsaturated Me-PGP-AR was lower in the vesicle fraction (11.1 ± 3%) (Supplementary Table 13). Phosphatidylglycerol-archaeol (PG-AR) was the second most abundant lipid in all fractions (18.8 ± 5.3%, 17 ± 3.7% and 32.6 ± 2.9% for cells, cell membranes and EVs respectively) and showed the highest degree of unsaturation (either 4 or 6 double bonds). The ratio of unsaturated to total PG-AR did not show a large variation between cellular (36.1 ± 4.1%), cell membrane (38.9 ± 6.4%) and extracellular fractions (31.3 ± 9%). Sulfated-diglycosyl-archaeol (S-2G-AR) showed relative abundances of 5.75 ± 4.4% (whole cells), 6.8 ± 3.9% (cell membrane) and 12.9 ± 1.5% (EVs) respectively, with negligible amounts of unsaturated lipids detected in the whole cell and cell membrane fraction.

**Figure 6.**
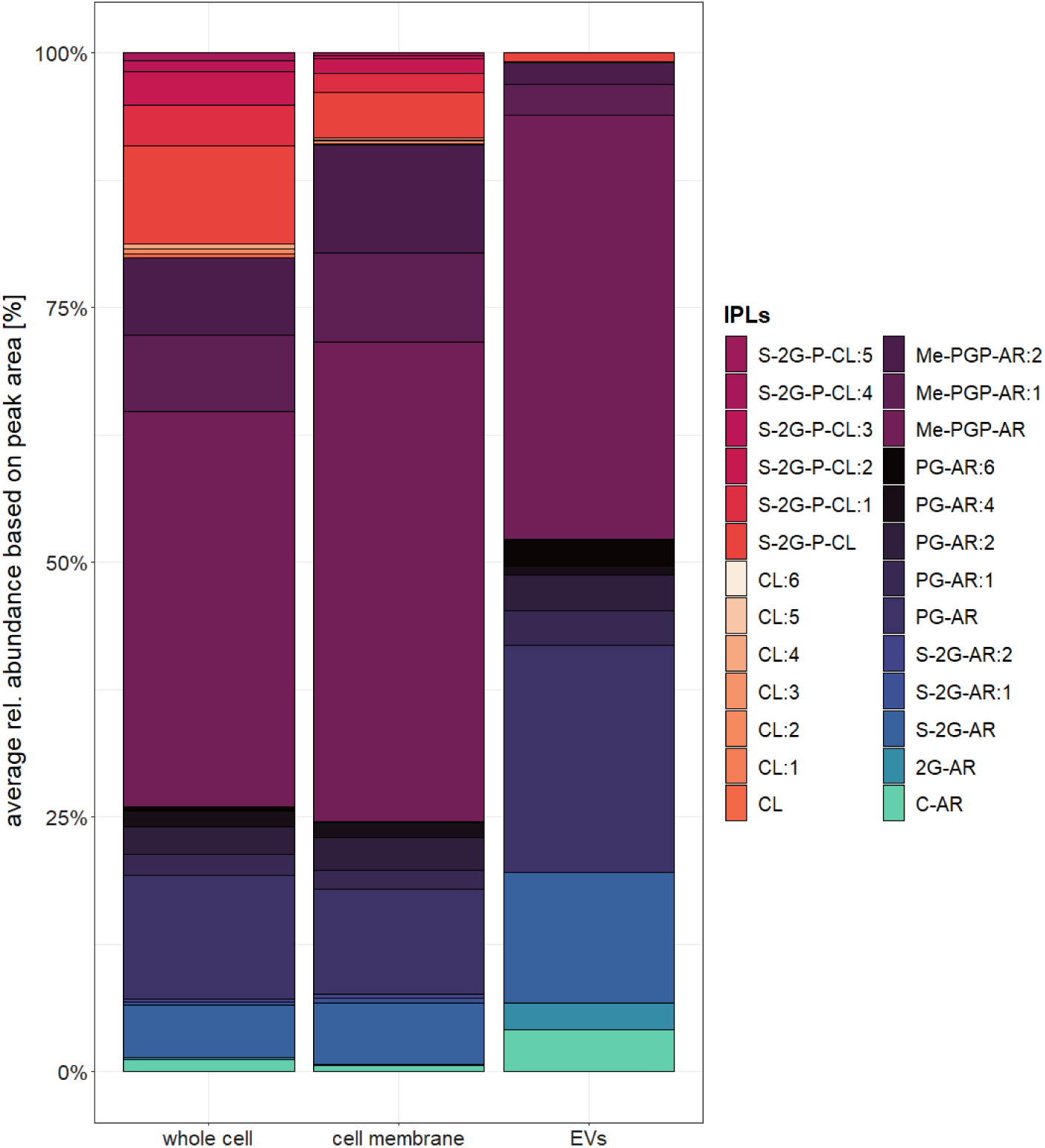
Average distribution of lipid compounds comparing whole cells, cell membranes and EVs of *H. volcanii*. The average (n = 3) relative abundance of lipids was calculated for each preparation; whole cells, cell membrane and extracellular vesicles (EVs) based on the peak area of the most abundant adduct for each compound. The distribution in the individual samples is shown in Supplementary Figure 17A. For the EV fraction, bands after density gradient purification were pooled together from 3 biological replicates. Lipids were identified based on their retention time, fractionation pattern and exact mass. *Compound abbreviations:* AR = archaeol (C_20_-C_20_ isoprenoidal chains), CL = cardiolipin, :nUS = lipid with n number of unsaturations, UK = unknown compound. Lipids with neutral headgroups: 1G = monoglycosyl, 2G = diglycosyl, C-AR = core-AR. Lipids with anionic headgroups: Me-PGP = phosphatidylglycerophosphate methyl esters, PG = phosphatidylglycerol, S-2G =sulfated diglycosyl, S-GP = sulfoglycophospho, 2P-GLY = diphosphoglycerol.

Lipids with a neutral headgroup, such as diglycosyl-archaeol (2G-AR) or no head group, such as core-archaeol (C-AR), were detected in all fractions but showed higher relative abundances in the EV samples (2.57 ± 0.19% and 4.1 ± 1%) compared to lipid extracts from cells and cell membranes (<1.2 ± 0.2%). A notable difference was also observed for dimeric phospholipids (or cardiolipins, CL). While they contributed 20.1 ± 9.7% and 9.04 ± 8.1% of the total lipids in whole cells and cell membrane samples respectively, they were almost undetectable in the EV samples (0.91 ± 0.41%). Interestingly, we were not able to detect any extended archaeol lipids (C_25_ instead of C_20_ isoprenoidal chains) with relevant concentrations in any of the samples, despite how common they are among many haloarchaea [45].

As expected, we could not detect any lipid compounds which were only present in the vesicular fraction but not in cells or cell membranes. However, the lipid composition of EVs differed significantly to that of cells and cell membranes when comparing the relative abundance patterns of different lipid groups. The distribution between unsaturated and saturated compounds shifts towards saturated lipids from 67.5 ± 2.7% and 68.7 ± 1.7% in whole cell and cell membrane extracts, to 84.4 ± 4.7% in EVs (Supplementary Table 13). In the vesicle fraction this is likely attributable to the absence of cardiolipins and the lower abundance of unsaturated Me-PGP-ARs.

### EV-associated RNA is taken up by H. volcanii cells

In order to test the ability for EVs to deliver the RNA cargo to a target organism, we used ^14^C-labelled uracil as a reporter to track the movement of RNA. EV preparations from the EV-deficient OapA knockout strain served as a control.

About 98% of the introduced radioactivity was taken up by both the parental strain and the OapA knockout strain over 6 days of growth. Subsequently, 1.90% of the radioactivity was detected in EV preparations of the parental strain, whereas only 0.11% was detected in EVs of the OapA knockout strain. After 20 min of incubation of the labelled EV preparations with fresh cells, we could detect a transfer of radioactivity into the unlabeled cells, with parental strain EVs transferring significantly more radioactivity than the OapA knockout strain EV preparation (Figure 7). Measurements after 90 min of incubation did not show a change of radioactive uptake from EVs of both strains, indicating that the transfer was already complete after 20 min. Thereby we confirm that the RNA enclosed in *H. volcanii* EVs can be internalized by *H. volcanii* cells in a short time frame.

**Figure 7.**
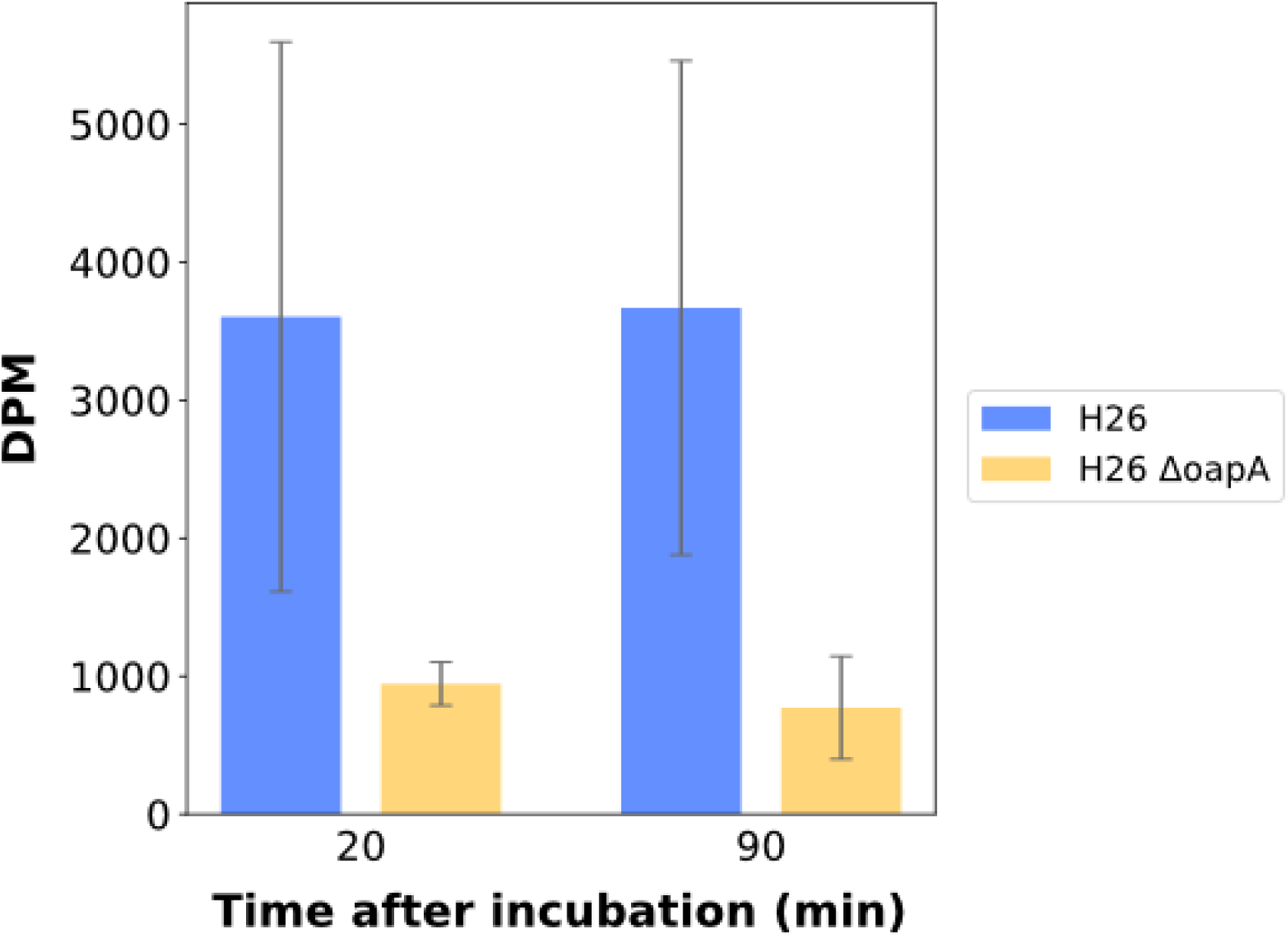
Transfer of radioactively labelled RNA by EVs. EVs were isolated from cells (H26 and H26 ΔoapA) incubated with radiolabeled uracil, resulting in EVs associated with radiolabeled RNA. EVs were then incubated with non-labelled cells and intracellular radioactivity in decays per minute (DPM) was measured 20 and 90 minutes post incubation using a scintillation counter.

## DISCUSSION

While more evidence arises that EVs play important roles in mediating important cellular functions in Bacteria and Eukaryota, there is still a disproportionate lack of information regarding the function and cargo of EVs in Archaea. Characterization of archaeal EV production and their biochemical composition can not only provide insights into the interactions between microorganisms in their unique environments, but also allow insights into the evolution of eukaryotic membrane trafficking mechanisms. EV production has been previously reported in haloarchaea [16], and here we used the haloarchaeal model organism, *H. volcanii*, to investigate the nature of these EVs and the mechanisms of EV production.

EV production by *H. volcanii* appeared to be influenced by temperature, while UV exposure and infection with a chronic virus had no significant influence under the conditions tested. EVs are heterogeneous in size, ranging from 50-150 nm. Analysis of the nucleic acid content of EVs produced by *H. volcanii*, as well as other haloarchaea, revealed that EVs are associated with RNA, as it has been described for some bacterial and eukaryotic EVs [46, 47], indicating that RNA associated EVs are conserved among all three domains of life. *Thermococcus onnurineus* (Euryarchaeida) has previously been reported to produce EVs containing RNA [41]; however, no characterization of EV-associated RNA was carried out for this organism. Treatment of EVs with nucleases did not eliminate the presence of EV-associated RNA. Additionally, TEM analysis comparing the wild type and a mutant with a destabilized S-layer, clearly showed that the surface of EVs is covered by the S-layer without any evidence for nucleic acids (Figure 1, Supplementary Figure 18C), and destabilization of the S-layer did not abolish the RNA cargo. Therefore, we infer that the RNA is internalized within EVs.

During preliminary data acquisition, we realized that a comprehensive picture of the EV composition is only gained when comparing the EV composition with the respective composition of the cell or the cell membrane, an approach that should be more common to studies investigating EVs.

While the RNA composition of *H. volcanii* EVs, both under normal growth conditions and under infection with a virus, appears to reflect intracellular levels to a certain extent, there is a distinct population of transcripts associated with EVs that does not correlate with the respective intracellular abundance, but are instead more enriched within EVs. The majority of highly enriched transcripts encode for tRNAs and rRNA, and we suggest that they are enriched due to both their structural stability and their high intracellular abundance. Both tRNAs and rRNAs have been observed at high abundancies in vesicle-associated transcriptomics in bacterial EVs [8, 25], and could therefore be a commonality among EVs from prokaryotic organisms. Interestingly, the most enriched mRNA that we detected was shown to be non-specifically fragmented in the EV-associated RNA fraction. Since we could not identify a common sequence or structural motif that would allow for a specific selection of particular RNAs to be enclosed into EVs, we suggest that the size, stability or both are a defining factor for packaging. Additionally, the positioning of an mRNA close to the cell envelope, such as the mRNA of the S-layer protein, could play a role in determining the RNA population of EVs. Results we obtained from EVs of viral infected cultures showed that the RNA composition did not change significantly upon infection in both cells and EVs; however, we detected viral RNAs in the cells and subsequently also in EVs, clearly representing the current transcriptional state of the EV-producing cell. When exposing cells to UV radiation, we subsequently observed changes to the RNA composition in EVs of UV-treated cells when compared to those of untreated cells (Supplementary Results, Supplementary Table 5). However, we did not acquire data to compare EV and intracellular transcriptomes, making it difficult to conclude with confidence whether these changes are due to changes in the cell. Nevertheless, since UV-treatment is known to influence the transcriptional landscape in *H. volcanii* cells [21], we assume that the changes observed in EVs are reflecting changes in the cell. In conclusion, we propose that RNA is taken up randomly into EVs, with transcripts that are highly enriched in the cell as well as transcripts that are translated at the cell envelope being preferably packaged. The respective cargo could be processed within EVs by RNases present in the vesicles (see protein content of EVs), leading to the degradation of mRNAs and a selection towards more stable RNAs (ncRNAs, tRNAs, rRNAs). Alternatively, there could also be a preselection for small-sized RNAs for packaging into EVs. Both scenarios lead to an RNA cargo representing a transcriptomic snapshot of the cell with a particular enrichment in RNAs with a regulatory potential (ncRNAs, tRNAs), as we observe in *H. volcanii* EVs.

The expression of ncRNAs in *H. volcanii* has been observed to shift dramatically under different conditions [48], and we predict that the population of packaged ncRNAs also reflects this shift. There are some notable, studied examples showing EV-packaged ncRNAs regulating gene expression in a receiving organism, such as EV-associated ncRNAs of *Vibrio fischeri* [8] and *Pseudomonas aeruginosa* [7], and we identify ncRNAs with regulatory potential associated with *H. volcanii* EVs. For example, we find a number of asRNAs overlapping with the start codon of various transposases that could potentially modulate the activity of transposases in a receiving organism. Unfortunately, the other identified ncRNAs do not have predicted functions. Currently, the nature of ncRNA-mediated regulation in Archaea is still unknown, with the majority of identified and predicted ncRNAs being uncharacterized. We have demonstrated that EVs of *H. volcanii* are able to transfer RNA between cells, and that RNA associated EVs are also produced by other haloarchaea. Therefore, we propose that halophilic archaea produce EVs as an intercellular communication mechanism to reflect the current intracellular state of the organism, and possibly influence gene expression in the receiving cell in response to environmental stimuli.

Proteomic analysis of EVs allowed us to draw conclusions about the mechanisms of the formation of EVs in haloarchaea. CetZ proteins were found particularly prominent in EVs of *H. volcanii* and *Hrr. lacusprofundi* [16]. However, EV production could still be observed from knockout strains of the enriched CetZ proteins (Supplementary Figure 17), suggesting that they do not play a significant role in EV formation in *H. volcanii*. CetZ proteins are known to be associated with the cell envelope [29], and we assume that this loose association could lead to enclosing of CetZ proteins during EV formation. Alternatively, the treatment that we used to prepare cell membranes could have also dissociated CetZ proteins from the membrane, leaving the impression that CetZ proteins are enriched in EVs. We also suggest that this could be true for other membrane-associated proteins.

In contrast, the knockout of a small GTPase (OapA), a Ras-superfamily GTPase with homology to a predicted Sar1/Arf1-GTPase that was also detected in PVs and EVs of *Hrr. lacusprofundi* [16], showed a very strong effect on EV formation. While the knockout of OapA results in a EV-deficient strain, overexpression of the small GTPase in the wild-type background leads to overvesiculation, further demonstrating the key role this protein plays in EV formation in haloarchaea. Rab and Arf GTPases, belonging to the Ras superfamily, are integral to the production of various vesicles in eukaryotic cells [49], including COP vesicles. COP vesicles regulate the trafficking of specialized lipids and proteins between the endoplasmic reticulum and Golgi apparatus [37]. The production of these vesicles requires the activation of the GTPase in order to recruit the coat complex, resulting in deformation of the cell envelope and subsequent budding of the vesicle [50, 51]. Deletion of this protein in Eukaryotes results in the elimination in the production of COP vesicles [52], and we have observed a similar suppression when knocking out the GTPase in *H. volcanii*, demonstrating that a functional Rab/Arf-related GTPase exists in Archaea and is able to regulate vesicle production. Homologous proteins of this new family of archaeal Ras-superfamily small GTPase can be identified across not only haloarchaea and Euryarchaeida, but also within other major branches in the archaeal domain, suggesting that this mechanism of EV production is widespread among specific clades of Archaea, specifically within Euryarchaeida and DPANN. Further, we observed that the archaeal GTPases group in accordance to their phylogeny, implying that these Archaea had acquired the gene early in their evolutionary past. Only a few phyla outside of DPANN and Euryarchaeida were found to contain this family of GTPase, forming a small branch containing *Asgardarchaeota, Thermoprotea, and Hydrothermarchaeota*. Within this branch, only specific clades of each phyla were represented, suggesting that the gene was attained in those clades through horizontal gene transfer events. Small GTPases within the Ras superfamily have been identified previously in Archaea, some of which clustering closely with known eukaryotic Ras GTPases [53]. These specific eukaryotic and eukaryotic-like GTPases contain a conserved aspartate in the G3 region that is also present in the *H. volcanii* GTPase, OapA, suggesting that the family of archaeal small GTPases identified in this study is also closely related to the eukaryotic Ras GTPases. Though homologs were not as widely identified among other clades, this could be due to the fact that some clades are represented by uncultured organisms and their respective MAGs or that the small GTPases present in those organisms are too divergent from the subfamily identified here. We opted for a more stringent search for homologous proteins across the archaeal domain that could have excluded other novel archaeal protein families more distantly related to the GTPase family identified here, but also still carry out the same function. For instance, only 3% of the surveyed *Asgardarchaeota* contained a homolog to OapA, yet they have been shown to contain small GTPases in close genomic proximity to other coatomer-like proteins, suggesting that they also contain a functionally similar GTPase [54]. Furthermore, we identified one ß-propeller repeat containing protein (WD40 domains) associated with EVs with homologs identified in EVs from *Hrr. lacusprofundi* [16]. Proteins with WD40 domains can also be identified in the coatomer of intracellular vesicles of Eukaryotes [37]. Therefore, we propose that proteins involved in EV formation in haloarchaea, along with other lineages of the archaeal domain, could represent evolutionary precursors to proteins facilitating intracellular vesicle formation in Eukaryotes. *Asgardarchaeota*, the currently known closest relatives to Eukaryotes [55], have been shown to encode homologs to ESCRT machinery proteins, a group of proteins that are involved in EV production in Eukaryotes [56] and have recently been shown to be crucial for EV formation in *Sulfolobus* [14]. *Asgardarchaeota* have also been shown to encode homologs to intracellular membrane trafficking proteins, such as Ras-like GTPases like those identified in this study, and WD40 domain proteins [54]. Provided that the Ras GTPases and ESCRT proteins in *Asgardarchaeota* function in a similar manner to those characterized for *Sulfolobus* and *H. volcanii*, this represents two major mechanisms of eukaryotic vesicle formation potentially combined within one archaeal organism similar to Eukaryotes. Finally, this new family of archaeal Ras-superfamily GTPases appears to be highly conserved among DPANN archaea, implicating that vesicle formation, or a related mechanisms involving the GTPase, could be very crucial for DPANN archaea that are known for their symbiotic lifestyle [57]. The identification of precursors of eukaryotic intracellular vesicle formation in both free-living and symbiotic Archaea could have implications for a revision of the eukaryogenesis hypothesis.

We found other proteins that could also play a role in EV function, such as those with enzymatic functions or transport related proteins. Enzymatic activity was detected for EVs from the abundant marine cyanobacterium, *Prochlorococcus* [58], suggesting that EV-associated proteins can facilitate specific reactions extracellularly. Components of ABC transport systems make up the overall majority of proteins associated with EVs of *H. volcanii*, and were also detected in high abundancies in EVs and PVs of *Hrr. lacusprofundi* [16] as well as other characterized EVs [14]. While this enrichment could be due to their high abundance in the cell envelope, the binding capacity of the EV-associated solute-binding proteins could also allow sequestration of rare nutrients that could be incorporated by the receiving cell [59]. Alternatively, EVs could play a role in the removal of obsolete proteins from the cell envelope, such as components of ABC transporters, allowing the cell to refresh the composition of the envelope to better adapt to their environment. Furthermore, we identified a highly enriched diadenylate-cyclase, an enzyme involved in the formation of cyclic di-AMP. These molecules are known secondary messengers in *H. volcanii* [38] and could be enriched with EVs, providing an additional mechanism of communication.

Analysis of the lipid composition of EVs in comparison to the lipid composition of whole cells and cell membranes revealed some unexpected differences. We were able to detect the major bilayer forming lipids PG-AR, Me-PGP-AR, S-2G-AR, C-AR, 2G-AR and cardiolipins, that were previously described for *H. volcanii* [60, 61] in all samples, albeit in different relative amounts. Me-PGP-AR and PG-AR were the two most abundant lipid species across all samples, while the cardiolipins (CL) contributed to a notable portion of the intact polar lipids (IPLs) in cells and cell membranes and were surprisingly only detected in low abundances in EVs. CLs are considered to be important for membrane curvature [62]; therefore, we expected them to be essential in EVs due to the high degree of bilayer curvature in the vesicles. However, Kellermann et al [60] observed that changing extracellular Mg^2+^ levels influence CL and Me-PGP ratios in *H. volcanii* and proposed that changes to the ratio of the two compounds are used to control membrane permeability in neutrophilic haloarchaea, in response to extracellular Mg^2+^ levels. As we cultivated *H. volcanii* in medium with a constant high Mg^2+^ concentration (174 mM) it is not surprising that Me-PGP-AR was the most prominent phospholipid species across all samples. This could also explain the absence of CLs in EVs, as Me-PGP-AR may be sufficient to ensure membrane stability in the smaller-sized EVs under high Mg^2+^ concentrations.

C-ARs and 2G-AR showed the opposite trend to cardiolipins, with an increase in their relative abundance in EVs compared to the cellular fraction. EVs of the hyperthermophilic *Sulfolobus solfataricus* were also shown to contain the same lipid species as the respective producing cells with significant shifts in the ratio of particular lipid compounds [63], similar to what we observe in *H. volcanii*. Differences between the lipid composition of cells and EVs could point towards a specific enrichment of particular lipid compounds in the EVs.

In summary, we show that EV production and the enclosing of RNA into EVs is common for multiple haloarchaeal species. We propose that the formation of EVs in haloarchaea is an active and conserved process, considering the conditionality of EV production along with their molecular composition that differs significantly from the originating cell, and the crucial involvement of a GTPase that is conserved among haloarchaea and other archaeal lineages. The enrichment of RNA with regulatory potential in EVs and the conservation of this process among different species lets us propose that halophilic Archaea utilize EVs as a communication mechanism influencing gene expression at a population-wide scale, as it has been proposed for some Bacteria [7, 8]. Finally, we propose that EV formation in haloarchaea, and potentially a wide range of other Archaea, is related to Ras superfamily GTPase-dependent intracellular vesicle trafficking in Eukaryotes. Together with vesicle formation by *Sulfolobus* species that is dependent on ESCRT-like proteins [14], and related extracellular trafficking facilitated by ESCRT proteins in Eukaryotes, archaeal EV production sheds light on the evolution of both intra- and extracellular vesicle trafficking in Eukaryotes and might help to elucidate the eukaryogenesis puzzle [64].

## Data availability

Raw data for resequencing of H26 Δ*oapA* mutant are available at ENA under project number PRJEB58368. Raw data for all RNA sequencing experiments for *H. volcanii* and *Hbt. salinarum* are available at ENA under project numbers PRJEB58342 and PRJEB58367 respectively. The mass spectrometry proteomics data have been deposited to the ProteomeXchange Consortium via the PRIDE [65] partner repository with the dataset identifier PXD038319 and 10.6019/PXD038319. Account details for reviewer: Reviewer account details: Username, Password: KcmThVJF. Lipid data www.pangea.de.

## Authors Contributions

J.M. performed the majority of the experimental work. L.J.G. and F.S. performed the lipid analysis. A.S. performed proteomics. Y.L. generated the AglB mutant. D.R.S. performed oapA phylogeny analysis. S.E. discovered the presence of RNA in EVs, conceived and led the study. I.G.D. and A.M. led the initial phase of the study. Some of the experiments were performed in A.M. laboratory. J.M. and S.E. performed the primary writing of the manuscript. All authors participated in the analysis and interpretation of the data and contributed to the writing of the manuscript.

## Acknowledgments

We thank the Max Planck-Genome-Centre Cologne (Cologne, Germany) for assistance with DNA and RNA sequencing. We thank Ingrid Kunze (MPI for Marine Microbiology, Bremen, Germany) for assistance with the experiments. We thank Thandi Schwarz (Ulm University) for help with the Northern Blots. We thank Jörg Soppa (Goethe-University Frankfurt) for providing the OapA knockout strain. We thank Thorsten Allers for providing the vector used in this study. We thank José Vicente Gomes-Filho (MPI for Terrestrial Microbiology, Marburg) for assistance with transcriptomic analysis. Finally, we want to thank the Max-Planck-Institute for Marine Microbiology and the Max-Planck-Society for continuous support.

## METHODS

### Strains and media

*Haloferax volcanii* strains and other haloarchaea used in this study are summarized in Supplementary Table 1. *H. volcanii* was either grown in minimal media HV-Ca supplemented with the SL10 trace elements and vitamins as described for DBCM2 [66] (HV-Cab), or nutrient rich media HV-YPC supplemented with the same trace elements and vitamins [66]. Cultures were grown in glass flasks aerobically at 120 rpm at the temperatures indicated. For auxotrophic strains that required additional nutrients, media was supplemented with uracil (50 µg/mL) and tryptophan (50 µg/mL), depending on the strain (Supplementary Table 1). *Hbt. salinarum* was grown in media described in [67], aerobically at 45 °C and 120 rpm. *Halorubrum lacusprofundi* was grown in DBCM2 media [66] aerobically at 28 °C (120 rpm). UV treatment was performed by pouring the culture into a petri dish that was then placed into a UV crosslinker from Biometra™ and exposed to 0.05 J of UV radiation. Infection of *H. volcanii* cultures with the virus, HFPV-1, was performed as described in [22], and infection was confirmed by PCR as previously described in [22].

### Generation of knock out strains

To construct plasmids for the deletion of *aglB* gene, PCR fragments of the upstream and downstream flanking sequences (∼ 530 bp) were amplified (primers listed in Supplementary Table 2). The upstream and downstream fragments were joined by Gibson assembly and the products were digested with BamHI and HindIII (restriction sites located on the outer primers) and ligated to pTA131 digested with BamHI and HindIII. The resulting plasmid was demethylated and then used to transform *H. volcanii* H26 using the two-step procedure (pop-in and pop-out) [68]. The expected mutations were verified by allele-specific PCR to detect the absence of the *aglB* gene on genome, yielding strain Δ*aglB*.

### Isolation and purification of EVs

For isolation of EVs from *H. volcanii*, cultures were grown at 45 °C in minimal media with serial dilution (two times in exponential growth to OD_600_ = 0.05) before being transferred into nutrient rich media and grown at 28 °C (unless otherwise specified). EVs were isolated and purified as described in [69]. Briefly, cells were removed at late stationary (∼ 144 hours growth) by centrifugation (4,500 x *g*, 40 min), and EVs were precipitated with the addition of polyethylene glycol (PEG) 6000 and incubation at 4 °C. EVs were subsequently pelleted by centrifugation (14,000 x *g*, 50 min, 4 °C) and after resuspending the pellet, remaining cell contaminations were removed by an additional centrifugation (14,000 x *g*, 10 min) and filtration (1 x 0.45 µm filter, 1 x 0.2 µm filter). Extracellular nucleic acids were removed with DNase I (New England Biolabs, 20 U/mL) and RNase A (New England Biolabs, 20 U/mL) [70]. The samples were further purified through an OptiPrep™ density gradient, yielding two bands containing EVs.

EVs from *H. lacusprofundi* were isolated and purified following methods in [16]. EVs from *Hbt. Salinarum* were isolated as described for *H. volcanii*, but temperature for synchronization and final growth were 45 °C in media described above.

### Transmission electron microscopy

Samples were adsorbed onto a copper grid (FCF200-Cu) for 3 min and negatively stained with 2% uranyl acetate for 45 s. Grids were imaged at 200 kV with JEOL JEM-2100 Plus transmission electron microscope.

### RNA extraction and transcriptomic analysis

RNA was extracted from cell pellets or EV pellets using TRIzol™ (Thermo Fischer Scientific). 1 mL TRIzol™ reagent was added to the pellet, homogenized by pipetting, and incubated at room temperature for 5 min. 0.2 mL chloroform was added to the sample, gently mixed via inversion, and incubated at room temperature for 10 min. The sample was then centrifuged at 4 °C for 10 min at maximal speed (∼20,000 x *g*). Upper phase was transferred to a new tube, and 500 mL isopropanol was added, mixed gently by inversion, and incubated at room temperature for 10 min. The sample was then centrifuged at 4 °C for 15 min at maximal speed. The supernatant was removed and pellets washed twice with ice-cold 75% ethanol. The remaining liquid was removed and the pellet was air-dried for 10 min. Pellets were resuspended in RNase/DNase free water.

Libraries (small RNA libraries) were prepared and sequenced at the Max Planck-Genome-Center (Cologne, Germany). Preliminary RNA sequencing experiments (see Supplementary Results) were conducted with one replicate, while the final RNA sequencing for both untreated and HFPV-1 infected *H. volcanii* were performed in triplicates. RNA sequencing for *H. salinarum* was conducted with one replicate of cellular RNA and two replicates of EV-associated RNA pooled together. Read mapping and calculations of gene expression and differential expression was performed using Geneious Prime® (2021.0.1). Reads were mapped to a compiled version of all genomic elements using the Geneious mapper (including a standard read trimming step) with 99% minimum overlap identity (90% minimum overlap identity for preliminary *H. volcanii* read mapping and *Hbt. salinarum* RNAseq). For samples with 3 or more replicates, differential expression was calculated with DESeq2. For samples with only one replicate, the default Geneious differential expression calculator was used. Transcripts were considered significant if transcripts per million (TPM) was greater than 10, log2 fold change was greater than 1, and p-value was lower than 0.05. Consensus sequences were predicted using MEME [71] with default settings. Sequence alignment and structural alignment among asRNA was predicted using locaRNA [72–74], with the temperature setting set to 28 °C.

### Northern blot

The Northern blotting protocol was adapted from [75]. Briefly, RNA was extracted as described above and separated on formaldehyde-MOPS agarose gels, with a final concentration of 2% formaldehyde and 2% NuSieve 3:1 agarose (Lonza). 5 µg RNA was denatured for 10 min at 70 °C with 1 X MOPS buffer (20 mM MOPS, 5 mM NaOAc, 1 mM EDTA, pH 7.0), 3.7% formaldehyde and loading dye (67 nM EDTA pH 8, bromophenol blue and xylene cyanol in deionized formamide). Samples were heat denatured for 10 min at 70 °C then placed on ice for 3 min before loading onto gel. The gel was run at 125 V for 3 to 4 hours and the RNA was then transferred to a Zeta-Probe GT membrane (Bio-Rad) by capillary action with 20 x SSC buffer (3 M NaCl, 300 mM Sodium Citrate, pH 7.0) and 2 x SSC buffer. The oligonucleotide probe is listed in Supplementary Table 2, and was labelled with [γ-^32^P] ATP using polynucleotide kinase (Thermo Fisher).

### Plasmid construction and expression of OapA

The coding region for *oapA* was amplified by PCR (primers listed in Supplementary Table 2), digested with *Nco*I and *Eco*RI (restriction sites located on primer extensions), and ligated into pTA1852 digested by *Pci*I and *Eco*RI. The plasmid, pTA1852 (provided by Thorsten Allers), is derived from pTA1392 [76] with a replacement of the 112 bp *Nde*I and *Not*I region containing an N-terminal 6 x His tag and a C-terminal 1 x StrepII tag with an N-terminal 7 x His tag and 2 x StrepII tag. Expression of tagged OapA (OapA_t_) on pTA1852 is controlled by tryptophan-inducible promoter, p.tnaA. The resulting plasmid was demethylated and transformed into *H. volcanii* strain H26, and plated onto HV-cab plates (1% agar).

Expression of OapA_t_ was adapted from [77]. Cultures were grown in HV-YPC supplemented with 200 µg/mL tryptophan at 28 °C until OD_600_ of approximately 1. Cultures were then supplemented with tryptophan by adding 10% of total volume 18% BSW containing 5 mg/mL tryptophan (final concentration of 450 µg/mL tryptophan) to the culture. Cultures were grown for 2 hrs at 28 °C before EVs were quantified as described below.

Affinity purification of OapA_t_ was modified from [78]. Cells from 500 mL culture were pelleted (11,000 x *g*, 40 min) and resuspended in 7 mL Binding Buffer (20 mM HEPES pH 7.5, 2 M NaCl, 1 mM PMSF). Cells were lysed by sonication (6 x 30 seconds at 35% amplitude) on ice, and treated with 20 µL DNase I (New England Biolabs, 20 U/mL) for 1 hr at 28 °C. Lysates were centrifuged (20,000 x *g*, 15 min, 4 °C) and filtered through 0.8 µm, 0.45 µm and 0.22 µm pore-size filters. The remaining flow through was incubated overnight with 1 mL Strep-Tactin® Sepharose® beads (iba-lifesciences) equilibrated with Binding Buffer and applied to a Poly-Prep chromatography column (Bio-Rad). Flow through was run twice on the column, and the column was then washed 5 x with Binding Buffer. The column was then incubated with 3 mL Elution Buffer (Binding buffer with 5 mM D-desthiobiotin) for 30 min, and flow through was concentrated using Vivaspin® 500 centrifugal concentrator (10,000 MWCO, Sartorius). Expression of OapA_t_ was then confirmed using Western blot (Supplementary Figure 15A).

### Protein extraction and analysis

Protein content of EVs was compared with protein content of cell membranes as described previously in Erdmann et al [16]. Proteins were isolated from purified EVs (triplicates of each upper band and lower band in density gradients) and host membranes (in triplicates) from untreated and UV-treated samples as described in [69], resulting in a total of 12 EV samples and 6 cell membrane samples. TCA precipitated protein mixtures were dissolved in 30 µL 1 x Laemmli sample buffer and loaded on Any kD™ Mini-PROTEAN® TGX™ Precast Protein Gels (Bio-Rad Laboratories, Germany). After short 3 cm SDS PAGE separation, the gels were visualized with Coomassie staining and each gel lanes cut into two slabs, which were processed individually. Proteins were in-gel reduced with dithiothreitol, alkylated with iodoacetamide and digested overnight with trypsin (Promega Mannheim, Germany). Resulting peptide mixtures were extracted twice by exchange of 5% of formic acid (FA) and acetonitrile, extracts pooled together and dried down in a vacuum centrifuge. Peptides were then re-suspended in 25 µL of 5% formic acid and a 5 µL aliquot was analyzed by LC-MS/MS on a nano-UPLC system Ultimate3000 series interfaced to a LTQ Orbitrap-Velos mass spectrometer (both Thermo Fisher Scientific, Bremen, Germany). The nano-UPLC was equipped with an Acclaim PepMap100 C18 75 µm i.d. x 20 mm trap column and a 75 µm x 15 cm analytical column (3 µm/100 Å, Thermo Fisher Scientific, Bremen, Germany). Peptides were separated using 80 min linear gradient; solvent A was 0.1% aqueous formic acid and solvent B was 0.1% formic acid in neat acetonitrile. Spectra were acquired using Data Dependent Acquisition (DDA) method and Top 20 approach; lock mass was set on *m/z* = 445.1200 (polydimethylcyclosiloxane). Three blank runs were performed after each sample analysis to avoid carryover. Acquired spectra were searched against *H. volcanii* proteins in NCBI database (June 2020, 12045 entries) by MaxQuant software (v. 1.6.10.43) using default settings and MBR (Matched Between Runs) option. False Discovery Rate (FDR) was 1%, variable modifications – methionine oxidized, cysteine carbamidomethylated and propionamide; two miscleavages allowed; minimal number of matched peptides – two. Relative quantification was performed using LFQ intensity values calculated by MaxQuant. Proteins were only considered present in EVs if matched with two or more peptides in all EV samples for that condition, and a respective LFQ value was identified in all EV samples for that condition.

Differential expression of proteins was calculated using R package, DEP (differential enrichment analysis of proteomics data) (v. 1.21.0) [79], based on the LFQ intensity values generated by MaxQuant. Differential expression between EV-associated proteins isolated from upper and lower bands of density gradient was calculated using three replicates of EVs from untreated cultures. For comparison of protein abundancies in EVs and cell membrane, upper and lower bands were pooled together for a total of 6 replicates for EV samples and 3 replicates of cell membrane samples. The threshold for significant enrichment in EVs was a log2 fold change greater than 1 and adjusted p-value lower than 0.05.

### Identification and phylogenetic analysis of Ras small GTPases across the archaeal domain

Homologs of small GTPase, HVO_3014, were identified in major archaeal clades using BLAST and analyzed with Interpro [31] to confirm the presence of signatures typical for P-loop GTPases and small GTPases belonging to the Ras superfamily. This resulted in 21 sequences from different organisms, including *H. volcanii* and *Hrr. Lacusprofundi*. These 21 sequences were used as a reference database in a DIAMOND [80] search using as query the entire protein content of a non-redundant set of 78,768 archaeal and bacterial genomes comprised of the genome taxonomy database (GTDB) species representatives (r207) [81] and the global catalog of earth’s microbiomes (GEM) OTU dataset [82] dereplicated at 95% average nucleotide identity using fastANI [83]. This search resulted in 96,121 hits, and 1,686 true positive hits were subsequently selected using an alignment score ratio approach [40]. This set was further manually curated, removing the only 5 bacterial GTPases based on protein phylogeny using FastTree 2 [84] and MUSCLE [85], as well as removing 15 sequences longer than 250 amino acids. This resulted in a final protein set of 1,666 archaeal GTPase sequences, representing the total diversity of this protein family in sequenced archaeal and bacterial genomes. The final dataset was aligned with MUSCLE and a phylogenetic tree was constructed using IQ-Tree [86] with ultrafast bootstrap analysis [87] using 1000 bootstrap replicates and default settings, auto-selecting the substitution model [88]. The phylogenetic tree was visualized on iTOL (v 6.6) [89] as unrooted, and taxonomy was mapped onto the resulting tree.

### Lipid extraction and analysis

For the total cell and cell membrane fraction, cells were harvested from 50 mL of *H. volcanii* liquid cultures in three biological replicates. The pellets were dissolved in 5 mL DBCM2 salt solution [66] and divided equally into two halves. For lipid analysis from total cells 2.5 mL of each replicate were divided into 300 µL aliquots. For extraction of cell membranes, 2.5 mL dissolved cell pellet from each replicate was diluted (1:1) with DBCM2 salt solution and sonicated with a microtip sonicator (MS 73 Sonoplus, Bandelin electronic, Germany) 3 times for 30 s on ice at 35% output. The lysate was treated with DNase I (30 min at 28°C, 10 µL per mL), and spun down in a table-top centrifuge (8,000 x *g*, 30 min, 4 °C) to remove cell debris. Cell membranes were pelleted by ultracentrifugation (MLS-5 rotor, 248,000 x *g* for 15 min). The pellets were dissolved in 1 mL DBCM2 salt solution and distributed into 4 x 250 µL aliquots. EVs from three biological replicates (200 mL cultures) were treated with DNase and RNase and purified with an Optiprep™ gradient (4 hr at 150,920 x *g*). The resulting EV bands were extracted separately from gradients. Instead of PEG 6000 precipitation, the samples were concentrated using filter columns (Vivaspin 6, 100,000 MWCO PES, Sartorius, Germany) at 4 °C with a maximal speed of 4,000 x *g* and washed twice with DBCM2 salt solution. Each gradient band was concentrated to 900 µL, from which 3 x 300 µL technical replicates were aliquoted. All samples were stored at -20 °C until lipid extraction.

For lipid extraction, samples were sonicated for 1 h in an ice-cooled ultrasonication bath and treated with a protocol based on [90]. Phase separation after the final centrifugation resulted in an upper lipid-containing organic phase, a lower metabolite-containing aqueous phase and a protein-containing pellet. The separate phases were isolated into combusted glass LC-MS vials, dried under constant N_2_ flow and stored at -20 °C until further analysis. Three 300 µL aliquots of sterile DBCM2 salt solution were treated with the same protocol as negative controls.

For ultrahigh performance liquid chromatography (UHPLC) coupled to mass spectrometry (MS) analysis, dried samples were resuspended in a solvent mixture of dichloromethane:methanol (1:9) and 0.4% of the total lipid extract were injected for total cells, cell membrane and negative control samples, 4% for EV samples from replicate 1 and 2% for all remaining EV samples. Measurements were performed on a Dionex Ultimate 3000 RS UHPLC system coupled to a maXis ultrahigh-resolution quadrupole time of flight tandem mass spectrometer (Q-TOF MS, Bruker Daltonics). Separation of archaeal lipids was achieved on a Waters Acquity UHPLC BEH C18 column (1.7 µm, 2.1 x 150 mm) at 65 °C using reverse phase chromatography from [91]. Briefly, a 26 min gradient was run at a flow rate of 400 µL min^-1^ beginning with 100% A (held for 2 min), followed by an increase to 15% B within 0.1 min and ramping to 85% B in 19 min, followed by 8 min re-equilibration with eluent B. Eluent A was MeOH:H_2_O (85:15) and eluent B was IPA:MeOH (50:50), both with addition of 0.04% HCO_2_H and 0.1% NH_3_. Analysis was performed in positive ionization mode, scanning from m/z 100 to 2000. MS2 scans were obtained in data dependent mode.

Output data were analyzed with the manufacturer’s software (DataAnalysis 4.4.2, Bruker Daltonics). Lipid compounds were identified based on retention time, fractionation pattern and exact masses [60, 92, 93]. Several technical replicates were measured for each sample type, of which representative replicates were selected for each biological replicate. Since the EV samples showed minimal differences in lipid distribution between bands after ultracentrifugation (Supplementary Figure 17B), gradient bands were pooled for each biological replicate for further analysis. The samples were compared with respect to their relative abundance distributions without absolute quantification as representative standards to correct for response factors for most of the detected compounds are lacking. While the fractions represented measurements of different percentages of their respective TLE, 0.4% for total cells and cell membrane and 4% for EV.1 and 2% for EV.2-3, we chose not to correct for dilution as it would not affect the relative abundance distributions.

The relative abundances were normalized per replicate and averages for each fraction were calculated from three biological replicates (total cells and cell membrane fraction) and from the upper bands after density gradient centrifugation from three biological replicates (EV fraction). Figures were created in R Statistical Software (v4.1.2; R Core Team 2021) with the ggplot2 [94], plyr [95] and dplyr packages [96].

### EV quantification

EVs were quantified from 2 mL of culture supernatant after removal of cells through centrifugation at room temperature (∼20,000 x *g*, 10 min twice, followed by 30 min). The supernatant was then filtered through a 0.22 µm pore filter. For quantification using fluorescence staining, MitoTracker® Green (Invitrogen) was added to the 2 mL (final concentration 500 nM), inverted to mix, and incubated at room temperature for 30 min. EVs were precipitated overnight at 4 °C using PEG 6000 (final concentration 10%) and pelleted by centrifugation (∼20,000 x *g*, 40 min, 4 °C). The EV pellet was resuspended in 200 µL 22% buffered sea water (BSW) [66], and transferred to a 0.5 mL microcentrifuge tube. Fluorescence was measured on a Spectrophotometer (DeNovix, DS-11 FX+) with blue excitation (470 nm) and emission between 514-567 nm. Base fluorescence of 22% BSW was subtracted from each value to remove background noise. For quantification using immunodetection, EVs were pelleted by centrifugation after PEG precipitation (20,000 x *g*, 4 °C, 40 min) and the supernatant was removed completely. The pellet was resuspended in 100 µL of 50 mM Tris-HCl, and stored at 4 °C until further use. 5 µL of the EV preparation was slowly spotted onto a nitrocellulose membrane (BioRad), and left to dry for 1.5 hours. After the membrane was dried, blocking was performed with blocking solution (60 g skimmed milk powder into 20 mL 1X TBS buffer [10X TBS buffer: 24 g/L Tris-HCl, 5.6 g/L Tris, and 88 g/L NaCl, with pH adjusted to pH 7.6 with HCl]) for 30 min, followed by incubation with the primary antibody (against HVO_2204, CetZ1, that was found to be highly enriched in EVs [16]) 1:1,000 diluted in blocking solution for 1 hr. The membrane was washed twice with 1X TBS-TT (10X TBS-TT is 10X TBS buffer with 5 mL/L Tween 20 and 5 mL/L Triton X) and once with 1X TBS before incubation with the secondary antibody (IgG anti-rabbit HRP conjugate, Promega) 1:1,000 diluted in blocking solution for 1 hr. Washing steps were repeated and chemiluminescence was visualized using Clarity Western ECL Substrate (Bio Rad). Relative chemiluminescence intensity was calculated using ImageJ [97]. Two different quantification methods were used, because each of them proved unsuitable for some conditions tested. We assume that enclosing CetZ1 into EVs can be influenced by particular conditions. Using CetZ1 [29] as a reporter gene for detection of EVs in culture supernatants (immunodetection) was unsuitable when testing temperatures dependencies (Supplementary Figure 2A and B) and did also not reflect results that we obtained for EVs from the *aglB* knockout strain (Supplementary Figure 16D). The fluorescence-based method proved unsuitable for quantification of EVs in virus infected cultures, because viral particles also appeared to be stained with the fluorescent dye (Supplementary Figure 2F). P-values are calculated by unpaired, two-tailed t-test.

### Tracking of EV uptake using 2-14C Uracil

To generate EVs containing radiolabeled RNA, uracil auxotrophic parental strain, H26 [68], and uracil auxotrophic deletion mutant Δ*oapA* were inoculated into 50 mL of HV-cab supplemented with a mix of unlabeled uracil and 14C labeled uracil (8.621 µg/mL final concentration, 25 µCi per culture) with an optical density (600 nm) of 0.05, each in triplicates. Cultures were grown at 28 °C for seven days before EVs were harvested. To harvest the EVs, cells were pelleted by centrifugation (4000 x *g*, 1 hr). The supernatant was filtered through a 0.22 µm pore filter to remove the remainder of larger contaminants. EVs in the flow through were then concentrated with Vivaspin® 20 (10,000 MWCO, Sartorius). The filters were washed three times with 22% BSW to remove residual unincorporated 14C uracil and subsequently concentrated to 500 µL of radiolabeled EVs per replicate. *H. volcanii DS2* was grown in HV-YPC media at 45 °C until OD (600 nm) of 1. 60 mL of culture were harvested by centrifugation (4000 x *g*, 20 min), washed with 6 mL HV-YPC and subsequently resuspended in 6 mL HV-YPC. For each replicate, 500 µL of cell concentrate were incubated with the 500 µL of EV concentrate in a heat block at 28 °C, 300 rpm. After 20 and 90 minutes post incubation, 300 µL were removed for measurement. The cells were pelleted (5 min, 10000 x *g*) and washed 3 times with 22% BSW to remove any residual EVs present. The resulting cell pellet was resuspended in 500 µL 22% BSW, added to 4 mL scintillation fluid (Ultima Gold™ XR, Perkin Elmer) and measured in a scintillation counter (Tri-Carb 4910 TR, Perkin Elmer).

## SUPPLEMENTAL INFORMATION

**Supplementary Results, Supplementary Figures 1-19 and Supplementary Tables 1-2 are provided as pdf file.**

**Supplementary Table 3: Differential expression calculated for transcripts from total vs small RNA libraries of EV associated RNA.** One replicate of RNA associated with EVs isolated from the upper band of a density gradient was sequenced using a total RNA library and a small RNA library. Read mapping (90% minimum overlap identity, TPM) and differential expression (log2 ratio) performed with Geneious™ (2021.0.1). (Excel file)

**Supplementary Table 4: Differential expression calculated for transcripts from EVs of upper versus lower band of a density gradient.** One replicate of RNA associated with EVs isolated from the upper band and the lower band of a density gradient. Read mapping (90% minimum overlap identity, TPM) and differential expression (log2 ratio) performed with Geneious™ (2021.0.1). (Excel file)

**Supplementary Table 5: Differential expression calculated for transcripts from EVs from untreated and UV treated cells.** One replicate of RNA associated with EVs isolated from untreated and UV treated cells. Read mapping (99% minimum overlap identity, TPM) and differential expression (log2 ratio) performed with Geneious™ (2021.0.1). (Excel file)

**Supplementary Table 6: Differential expression calculated for EV associated transcript normalized with intracellular levels**. RNA was extracted from purified EVs and the respective cells in triplicates. Read mapping (99% minimum overlap identity, TPM) and differential expression (log2 ratio, p-value) were calculated with DESeq2 in Geneious™ (2021.0.1). (Excel file)

**Supplementary Table 7: ncRNA enriched in EVs.** List of the ncRNA that are highly expressed and with a high fold change in *H. volcanii* EVs. Data taken from Supplementary Table 6. (Excel file)

**Supplementary Table 8: Differential expression calculated for transcripts from *Hbt. salinarum* EVs normalized to intracellular levels.** RNA was extracted from purified EVs (duplicates) and the respective cells (one replicate). Read mapping (90% minimum overlap identity, TPM) and differential expression (log2 ratio) performed with Geneious™ (2021.0.1). (Excel file)

**Supplementary Table 9: Proteins enriched in EVs after normalization with the protein content of cell membranes.** Protein content of EVs was pooled from upper and lower bands in three replicates (total of 6 EV replicates) and quantities were compared with three replicates from host cell membrane preparations. Quantity was estimated using MaxQuant (v. 1.6.10.43) and differential expression analysis (log2 fold change, adjusted p-value) was calculated with DEP (v. 1.21.0) [79]. (Excel file)

**Supplementary Table 10: Proteins enriched in EVs from UV-treated cells after normalization with the protein content of respective cell membranes.** Protein content of EVs from UV treated cells was pooled from upper and lower bands in three replicates (total of 6 EV replicates) and quantities were compared with three replicates from respective host cell membrane preparations. Quantity was estimated using MaxQuant (v. 1.6.10.43) and differential expression analysis (log2 fold change, adjusted p-value) was calculated with DEP (v. 1.21.0) [79]. (Excel file)

**Supplementary Table 11: Proteins identified as present in all EV samples.** Protein content of EVs from untreated cells (3 replicates from upper and lower bands of density gradient each) and UV treated cells (3 replicates from upper and lower bands of density gradient each) was pooled (total of 12 EV replicates). Label-free quantities (LFQ) were calculated using MaxQuant (v. 1.6.10.43) and averaged over all replicates. Proteins were only considered present if peptide count was greater than or equal to 2 in all replicates and all replicates had a corresponding LFQ value. (Excel file)

**Supplementary Table 12: Taxonomy of Archaea identified to contain Ras-superfamily small GTPase.** 1,666 archaeal organisms out of 78,738 archaeal and bacterial genomes were identified to contain a homolog to Ras-superfamily small GTPase, OapA (see Methods). Taxonomy listed according to genome taxonomy database release (r207). (Excel file)

**Supplementary Table 13: Mass spectrometry peak areas and relative abundances of lipid compounds extracted from whole cells, cell membranes and EVs of *H. volcanii*.** Intact polar lipids were extracted from whole cells, cell membranes and vesicles of *H. volcanii* and measured with a Q-TOF MS (Bruker Daltonics). Output data were analyzed with the manufacturer’s software (DataAnalysis 4.4.2, Bruker Daltonics) and lipid compounds were identified based on retention time, fractionation pattern and exact masses and quantified via mass spectrometry peak area. (Excel file)

