## Supplemental Material for "Extracellular vesicles of Euryarchaeida: precursor to eukaryotic membrane trafficking"

#### Supplementary Results

***EV-associated RNA is best analyzed when using small RNA libraries and normalizing EV RNA content with host cell RNA content***

To determine the nature of the EV enclosed RNA, total RNA and small RNA (enriching for transcripts below 150 nt in length) libraries were prepared from EV-extracted RNA. When comparing sequencing results from both libraries, we observed a drastically different transcriptional profile (Supplementary Table 3). Around 95% of reads from total RNAseq mapped to ribosomal RNA (rRNA) and only 17 transcripts recruited enough reads to reach the threshold (TPM > 10). In contrast, the small RNA library revealed a more diverse array of transcripts with transfer RNA (tRNA) being the most dominant RNA species (around 85% of reads) and 264 transcripts identified within the threshold. Further, over 2000 transcripts were only identified in the small RNA library and not in the total RNA library, the majority of them being tRNAs and non-coding RNAs (ncRNA), indicating that the total RNA library excludes important smaller transcripts. Therefore, we decided to use small RNA libraries for further analysis of EV-associated RNAs, as this seems to yield a more accurate picture of the RNA composition of EVs.

We also compared transcripts from EVs in the upper and lower bands in density gradients to determine whether the bands represented different subpopulations of EVs with respect to RNA content (Supplementary Table 4). Indeed, we identified transcripts that were only present in the upper band (app. 200) or only in the lower band (80). However, the abundance of these transcripts was below the threshold (TPM > 10) and they were disregarded. Overall, the RNA composition between both bands was mostly identical, with few outliers. We concluded that the RNA composition alone is not the differentiating factor between the two subpopulations of EVs in different density gradient bands, and pooling the bands for further sequencing analysis is acceptable.

Since vesicle production could be linked to UV exposure [1], we aimed to determine whether subjecting cultures to UV radiation would alter the RNA composition of EVs. Indeed, 145 transcripts appeared to be present in a higher abundance (log2 fold change > 1) in the UV-treated sample, and 32 transcripts were present in a higher abundance (log 2 fold change < -1) in the untreated sample (Supplementary Table 5). This population of EV-associated RNA from UV-treated cultures included all forms of RNA, including mRNAs, tRNAs, rRNAs and ncRNAs. Nevertheless, we realized that without determining transcriptional changes within the cells, it is difficult to distinguish transcripts that are associated with EVs as a response to UV exposure and transcripts that are present in EVs simply due to changes in intracellular levels. In order to differentiate between random packaging and potentially selective packaging of RNA into EVs, it became clear that sequencing intracellular RNA at the time of EV harvesting was imperative for any kind of analysis.

#### Supplementary Figures

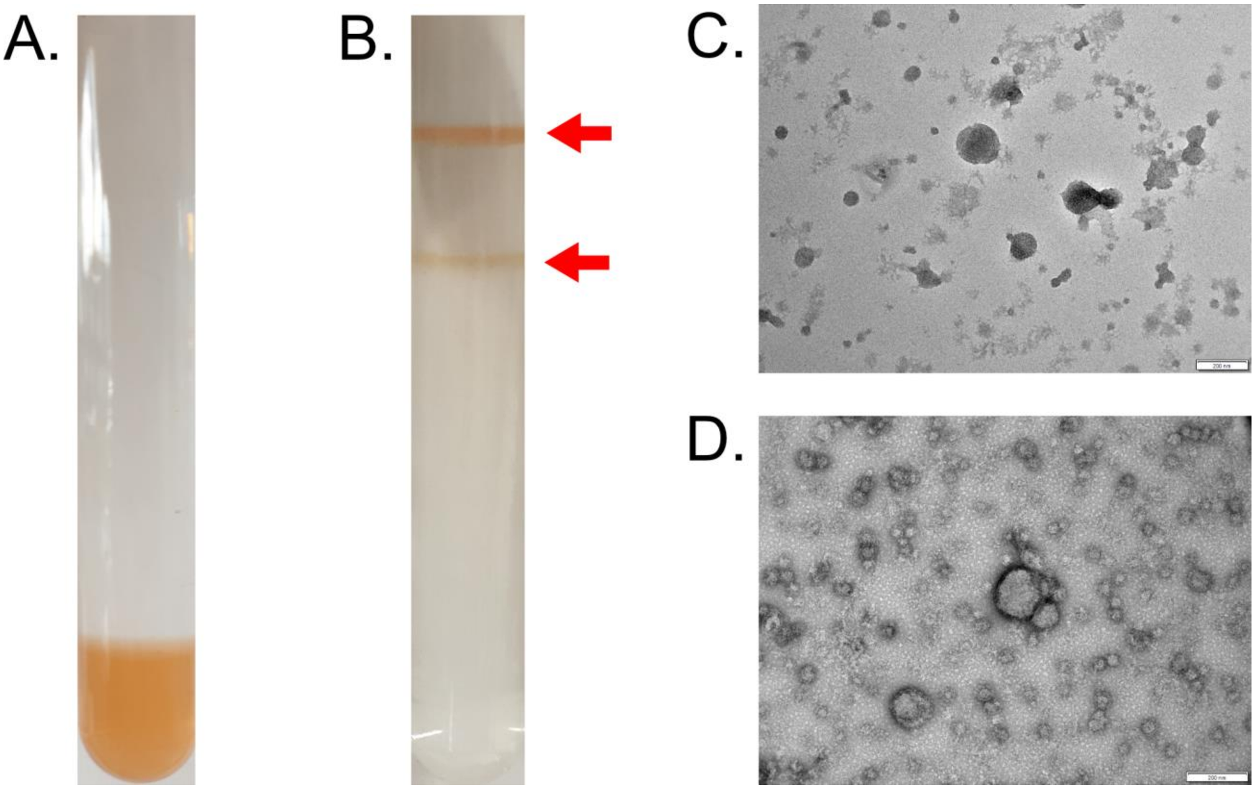


**Supplementary Figure 1. Purification of *H. volcanii* H26 EVs by Optiprep™ density gradient purification.** Gradient before **(A)** and after **(B)** ultracentrifugation. Red arrows indicate upper and lower band. Transmission electron micrograph of EVs isolated from upper **(C)** and lower **(D)** bands. Size bar: 200 nm.

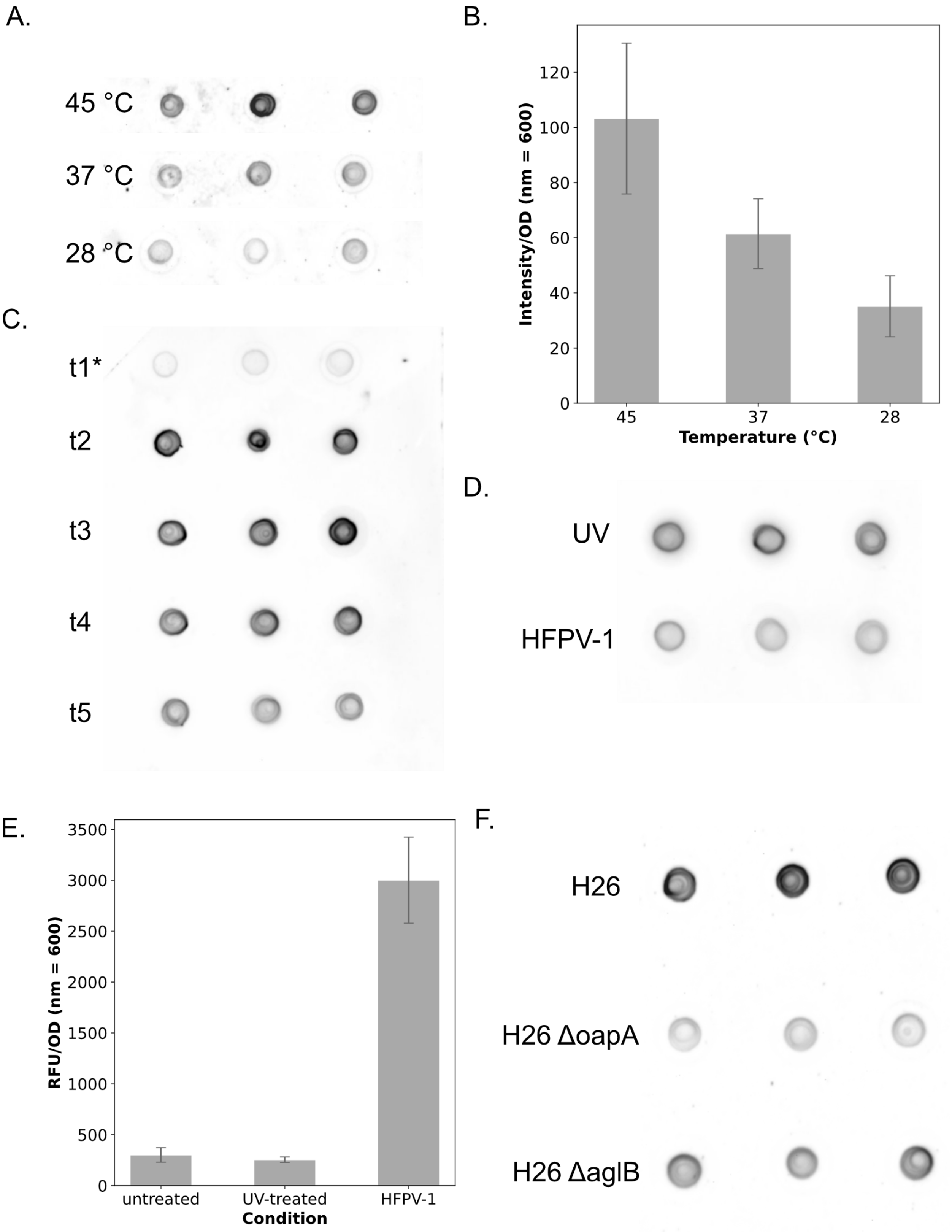
A.
45 °C
37 °C
28 °C
B.
Intensity/OD (nm = 600)
0
20
40
60
80
100
120
45
37
28
Temperature (°C)
C.
t1*
t2
t3
t4
t5
D.
UV
HFPV-1
E.
RFU/OD (nm = 600)
0
500
1000
1500
2000
2500
3000
3500
untreated
UV-treated
HFPV-1
Condition
F.
H26
H26 ΔoapA
H26 ΔaglB

**Supplementary Figure 2. EV quantification of *H. volcanii* cultures grown under different conditions, through growth and gene knockouts.** **(A)** Spot blot for quantification of EVs in *H.volcanii* culture supernatants grown at 45, 37 and 28 °C. **(B)** Bar plot representing spot blot **(A)**. **(C)** Spot blot for quantification of *H. volcanii* grown at 28 °C with time points taken at 45.3, 68.5, 94, 118.3, and 140.5 hours. Asterisk represents sample diluted by a factor of 2. **(D)** Spot blot for quantification of *H. volcanii* grown with either UV or viral stress. **(E)** Bar plot representing fluorescence-based quantification of EVs of *H. volcanii* grown with either UV or viral stress. **(F)** Spot blot for quantification of *H. volcanii* H26 background strains with knockouts of OapA and AglB.

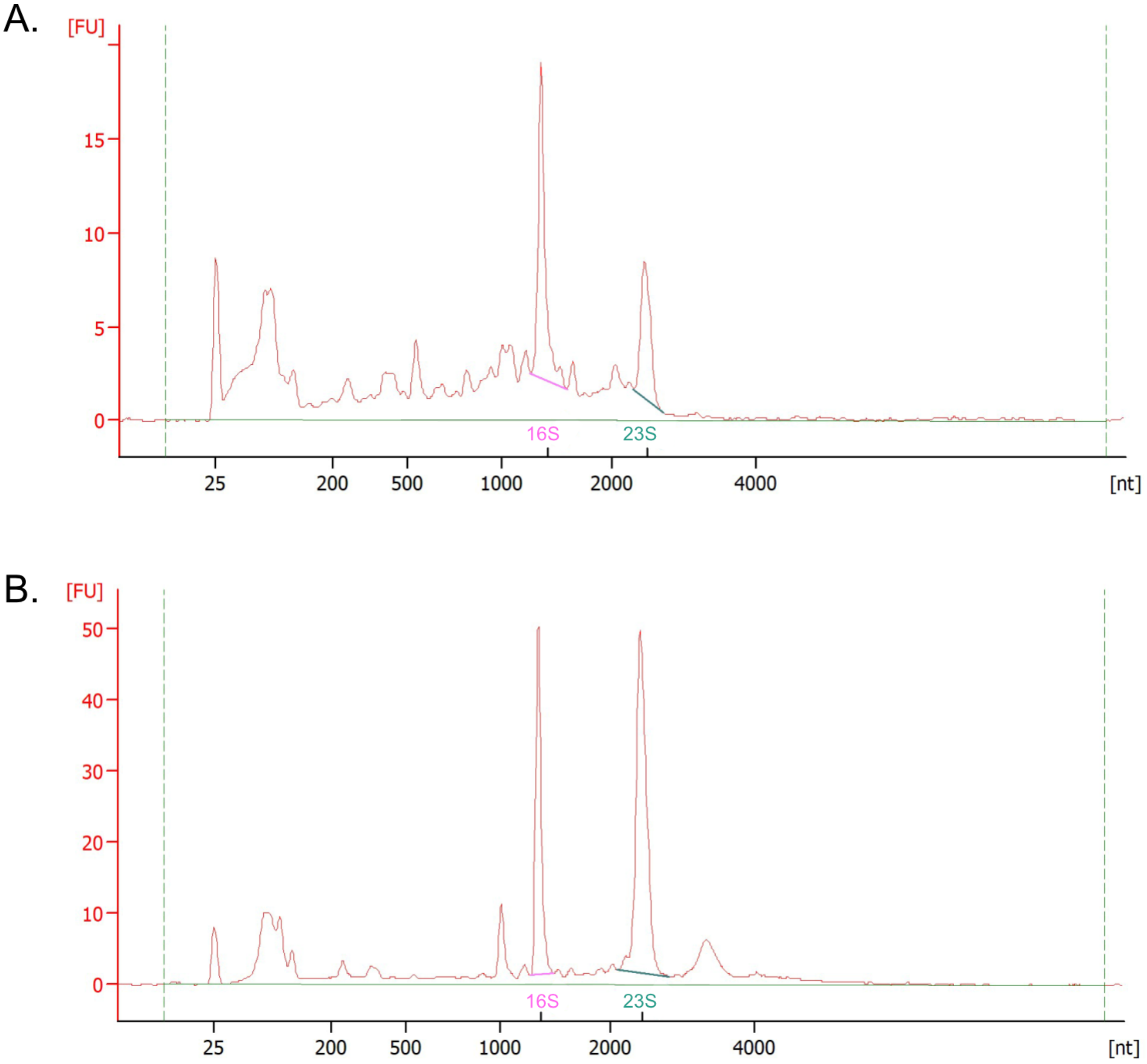


**Supplementary Figure 3. Electropherograms of EV-associated RNA (A) and cellular RNA (B).** Capillary electrophoresis demonstrates differences in size distribution between RNA isolated from EVs and cells of *H. volcanii*.

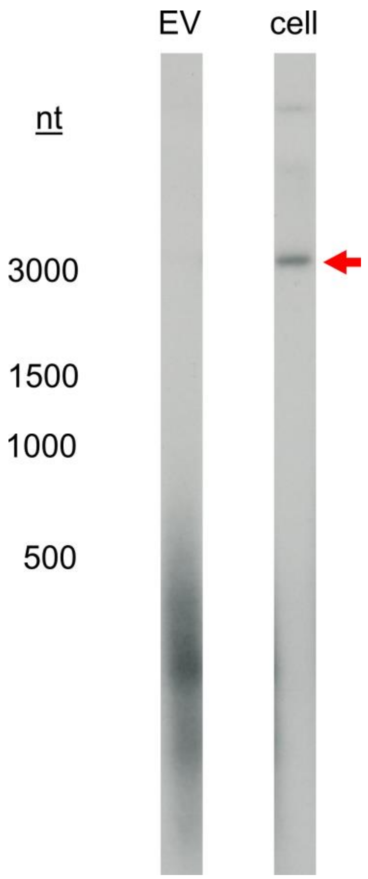


**Supplementary Figure 4. Northern blot with EV and cellular RNA probed for HVO_2072.** Red arrow indicates full-length transcript. Northern was conducted in duplicates, but only one replicated is presented here.

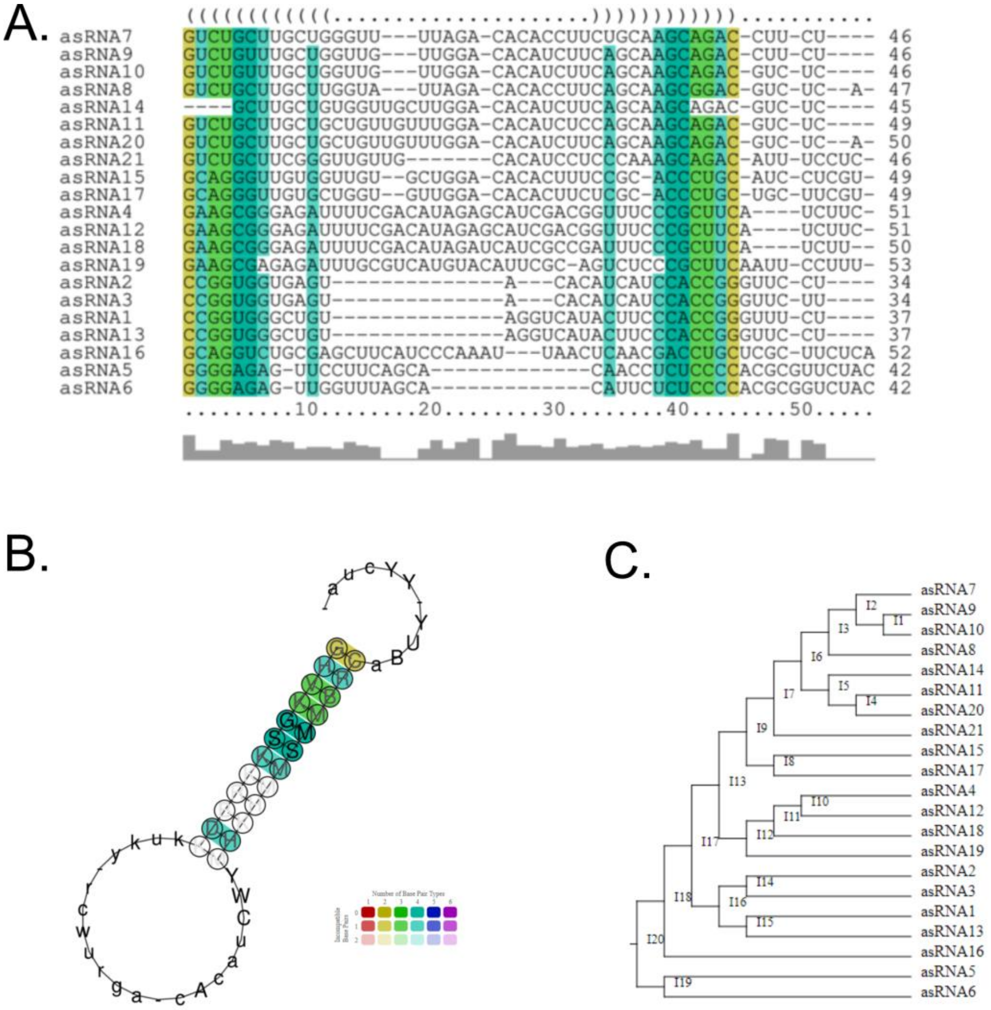


**Supplementary Figure 5. Secondary structure alignment prediction of EV-associated asRNA from LocARNA [2–4]. (A)** Sequence alignment with RNAalifold consensus structure. **(B)** Predicted consensus secondary structure with legend. **(C)** Hierarchical clustering based on similarities of sequences.

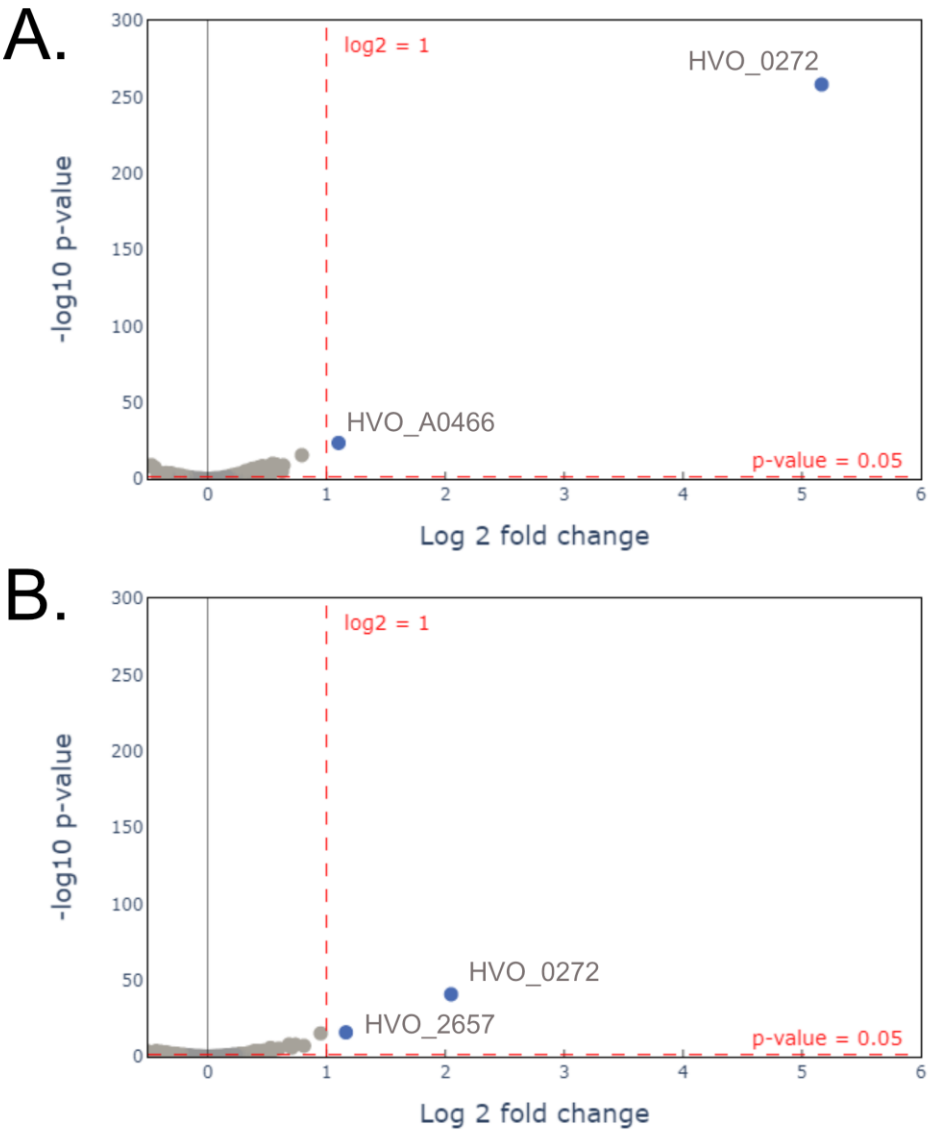


**Supplementary Figure 6. Volcano plots of intracellular and EV-associated transcripts comparing cultures infected with HFPV-1 and uninfected control cultures. (A)** Differential expression of transcripts in EVs from infected versus uninfected cultures. **(B)** Differential expression of intracellular transcripts from infected versus uninfected cultures. Cells and EVs were isolated at late stationary phase of growth. Volcano plots only depict transcripts that had an average TPM greater than 10 in either infected or uninfected samples.

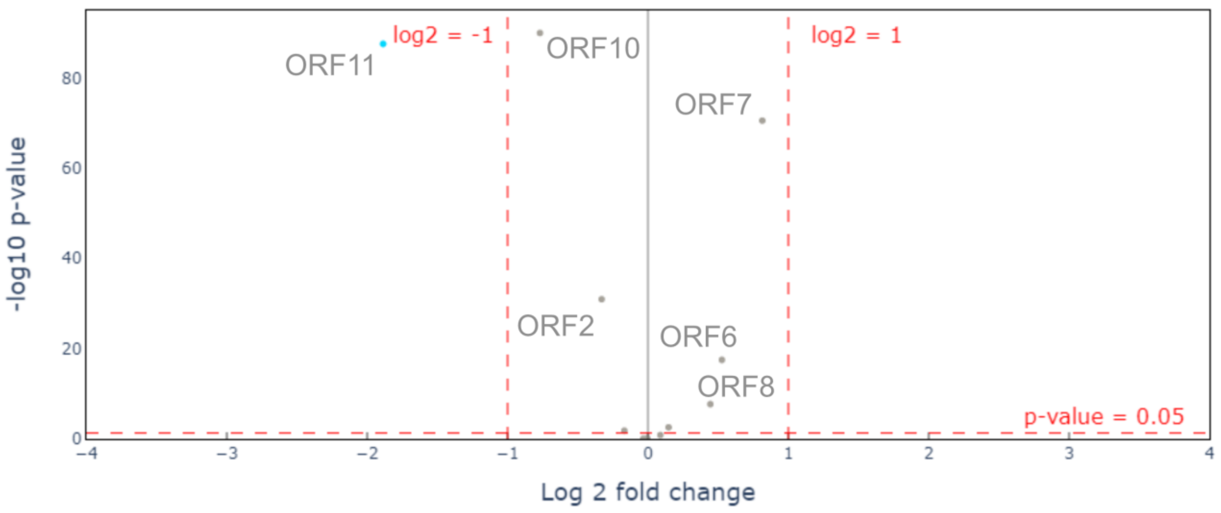


**Supplementary Figure 7. Volcano plot comparing viral transcript abundance between EV-associated RNA and cellular RNA.** RNA isolated from EVs and cells of cultures infected with HFPV-1 during late stationary phase.

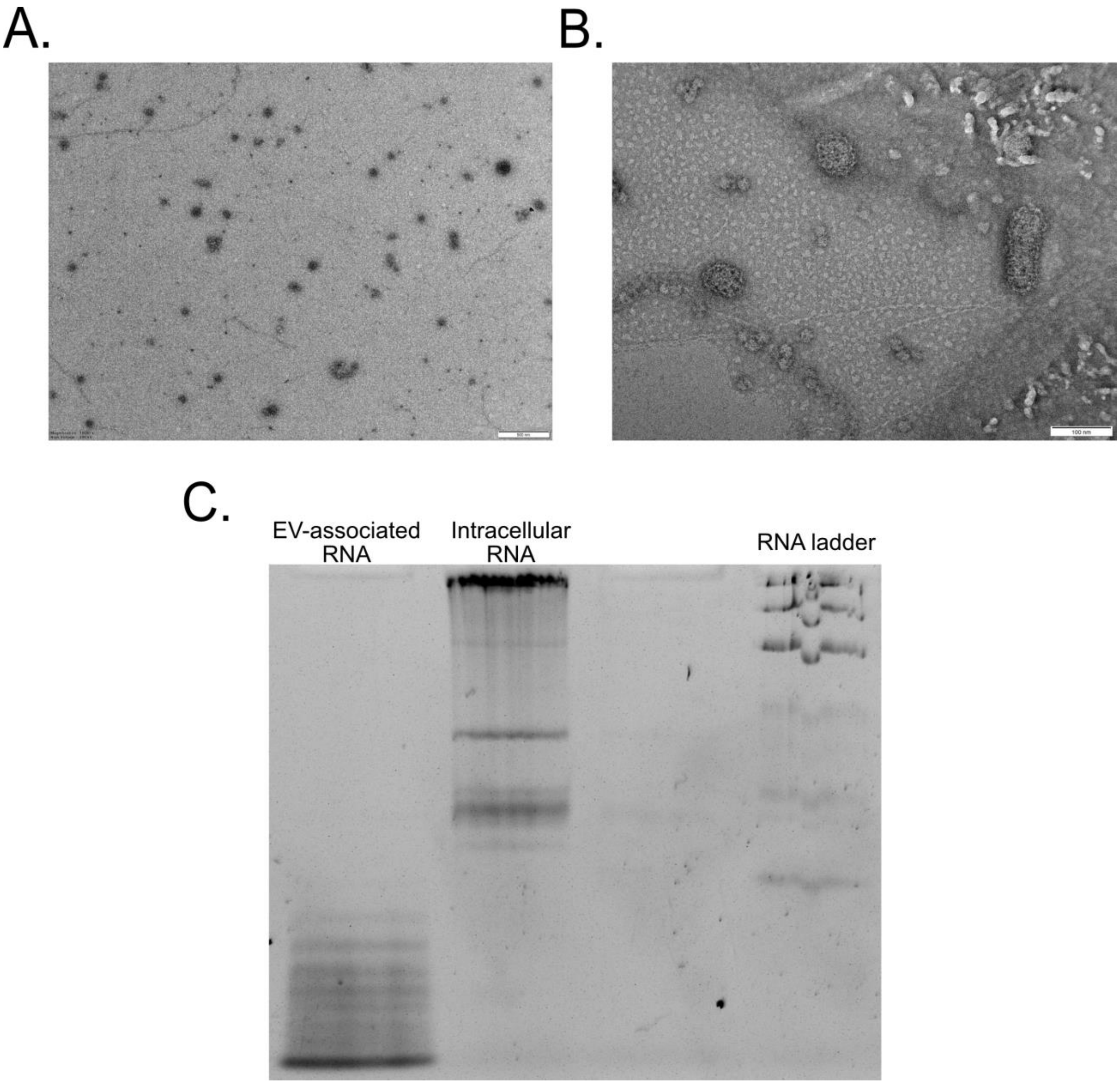


**Supplementary Figure 8. EVs from other haloarchaea. (A)** Transmission electron micrograph of purified EV from *Hbt. salinarum.* Scale bar: 500 nm. **(B)** Transmission electron micrograph of purified EVs from *Hrr. lacusprofundi.* Scale bar = 100 nm. **(C)** RNA extracted from gradient purified EVs and cells of *Hrr. lacusprofundi* on a 12.5% Urea-PAGE gel.

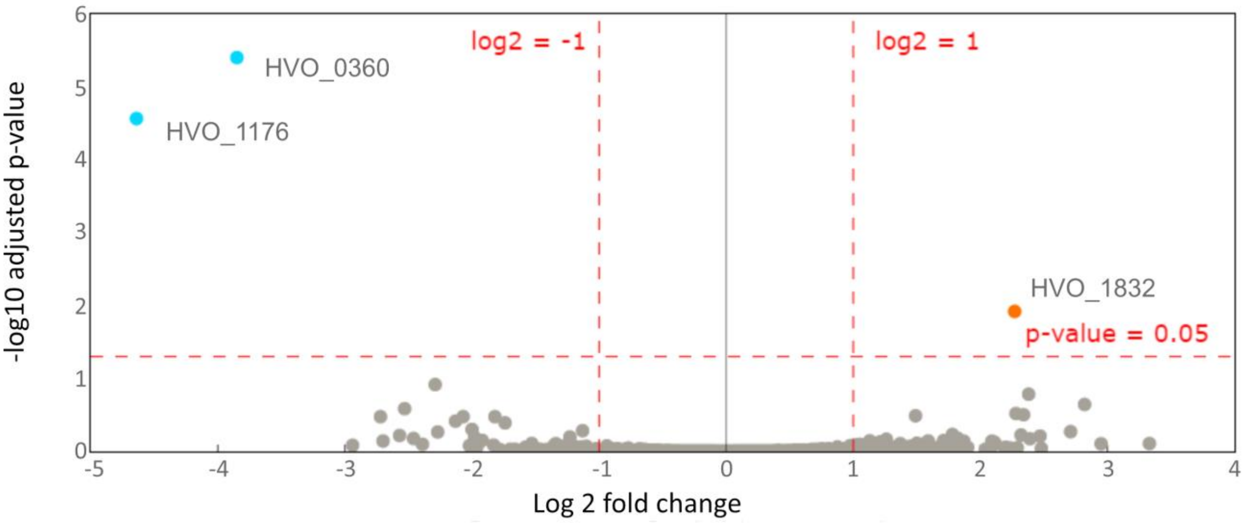


**Supplementary Figure 9. Volcano plot comparing protein content from *H. volcanii* EVs isolated from upper and lower bands of Optiprep™ density gradient.** EVs were isolated from three replicates at stationary phase of growth.

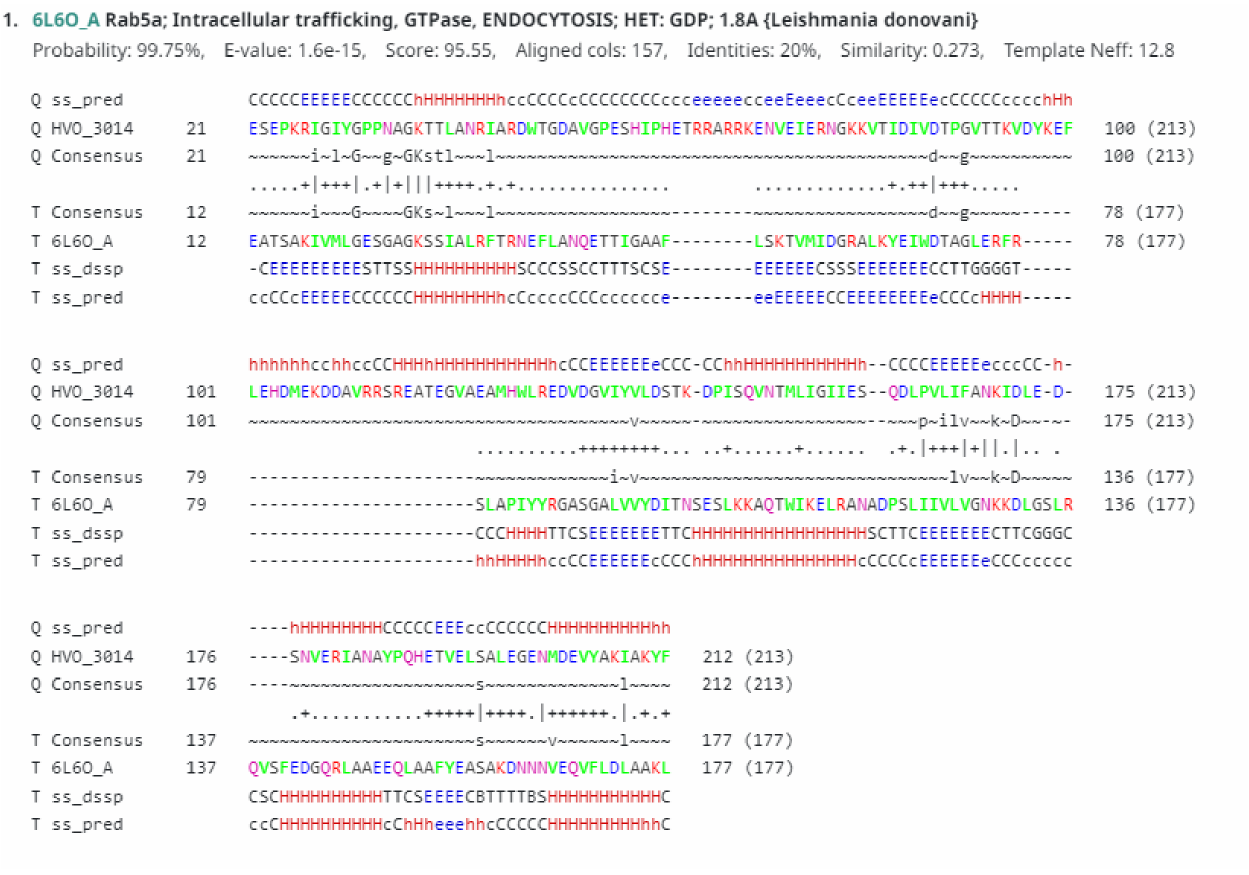

1. 6L6O_A Rab5a; Intracellular trafficking, GTPase, ENDOCYTOSIS; HET: GDP; 1.8A {Leishmania donovani}

Probability: 99.75%, E-value: 1.6e-15, Score: 95.55, Aligned cols: 157, Identities: 20%, Similarity: 0.273, Template Neff: 12.8

```
Q ss_pred                CCCCCEEEEECCCCCChHHHHHHHhccCCCCcCCCCCCCCccceeeeecceeEeeecCceeEEEEEecCCCCCccccchHh
Q HVO_3014      21      ESEPKRIGIYGPPNAGKTTLANRIARDWTGDAVGPESHIPHETRRARRKENVEIERNGKKVTIDIVDTPGVTTKVDYKEF   100 (213)
Q Consensus     21      ~~~~~~i~l~G~~g~GKstl~~~l~~~~~~~~~~~~~~~~~~~~~~~~~~~~~~~~~~~~~~~~~~d~~g~~~~~~~~~~~~   100 (213)
                        .....+|+++|.+|+|||++++.+.+...............        .............+.++|+++.....
T Consensus     12      ~~~~~~i~~~G~~~~GKs~l~~~l~~~~~~~~~~~~~~~~~~~--------~~~~~~~~~~~~~~~~~d~~g~~~~~-----   78 (177)
T 6L6O_A       12      EATSAKIVMLGESGAGKSSIALRFTRNEFLANQETTIGAAF--------LSKTVMIDGRALKYEIWDTAGLERFR-----   78 (177)
T ss_dssp               -CEEEEEEEEESTTSSHHHHHHHHHHSCCCSSCCTTTSCSE--------EEEEEECSSSEEEEEEECCTTGGGGT-----
T ss_pred                ccCCcEEEEECCCCCCHHHHHHHHhcCccccCCCcccccce--------eeEEEEECCEEEEEEEEeCCCcHHHH-----

Q ss_pred                hhhhhhcchhccCCHHHhHHHHHHHHHHHhcCCEEEEEEeCCC-CChhHHHHHHHHHHHh--CCCCEEEEEecccCC-h-
Q HVO_3014      101     LEHDMEKDDAVRRSREATEGVAEAMHWLREDVDGVIYVLDSTK-DPISQVNTMLIGIIES--QDLPVLIFANKIDLE-D-   175 (213)
Q Consensus     101     ~~~~~~~~~~~~~~~~~~~~~~~~~~~~~~~~~~~~~~~v~~~~~-~~~~~~~~~~~~~~~~~~~~--~~~p~ilv~~k~D~~-~-   175 (213)
                                              ..........++++++++... ..+......+......  .+.|+++|+||.|.. .
T Consensus     79      ----------------------~~~~~~~~~~~~~~~i~v~~~~~~~~~~~~~~~~~~~~~~~~~~~~~~~~~~lv~~k~D~~~~~~   136 (177)
T 6L6O_A       79      ----------------------SLAPIYYRGASGALVVYDITNSESLKKAQTWIKELRANADPSLIIVLVGNKKDLGSLR   136 (177)
T ss_dssp               ----------------------CCCHHHHTTCSEEEEEEETTCHHHHHHHHHHHHHHHHHSCTTCEEEEEEECTTCGGGC
T ss_pred                ----------------------hhHHHHhccCCEEEEEEcCCChHHHHHHHHHHHHHHHcCCCCcEEEEEEeCCCccccc

Q ss_pred                ----hHHHHHHHHCCCCCEEEccCCCCCCHHHHHHHHHHhh
Q HVO_3014      176     ----SNVERIANAYPQHETVELSALEGENMDEVYAKIAKYF   212 (213)
Q Consensus     176     ----~~~~~~~~~~~~~~~~~~~~~~~~~~s~~~~~~~~~~~~~~~~~~~~~l~~~~~   212 (213)
                            .+...........+++++|++++.|++++++.|.+.+
T Consensus     137     ~~~~~~~~~~~~~~~~~~~~~~~~~~~~~~~~~~~~~~s~~~~~~~v~~~~~~~l~~~~   177 (177)
T 6L6O_A       137     QVSFEDGQRLAAEEQLAAFYEASAKDNNNVEQVFLDLAAKL   177 (177)
T ss_dssp               CSCHHHHHHHHHHTTCSEEEECBTTTTBSHHHHHHHHHHHC
T ss_pred                ccCHHHHHHHHHHcChHheeehhcCCCCCHHHHHHHHHhhC
```

**Supplementary Figure 10. HHpred alignment of HVO_3014 with Rab5a GTPase from *Leishmania donovani*.** The protein sequence of HVO_3014 was analyzed using HHpred [1, 5] to identify domain-based homologous proteins using the default settings. Alignment with the best match to endosomal sorting protein, Rab5a GTPase, from unicellular Eukaryote, *Leishmania donovani*, is shown here.

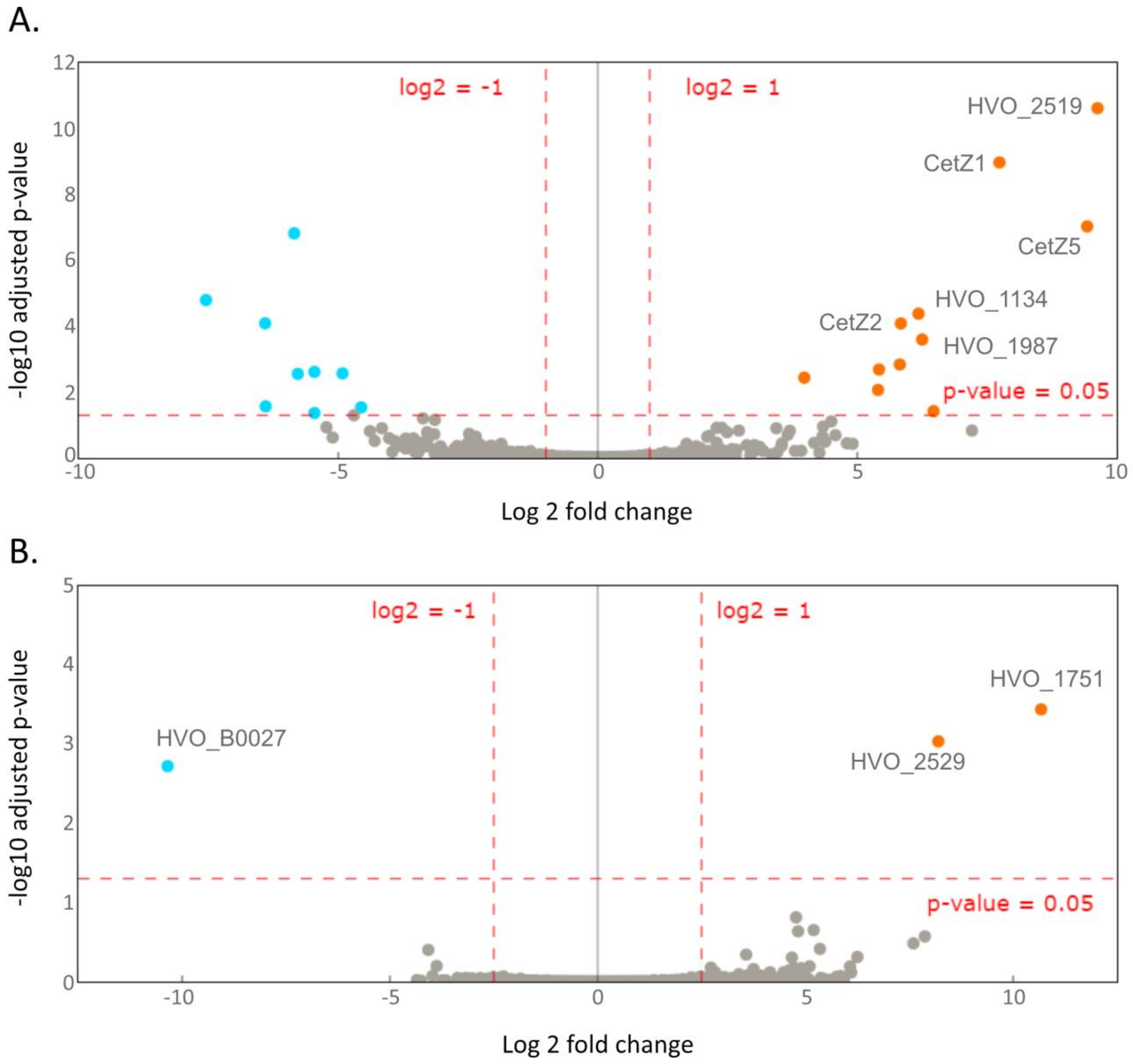


**Supplementary Figure 11. Volcano plots depicting differences in protein abundance from UV-treated cultures. (A)** Proteins isolated from EVs of UV-treated cultures are compared to their respective cell membrane protein content. **(B)** EV-associated proteins from UV-treated cultures are compared to EV-associated proteins from untreated cultures. Raw data found in Supplementary Table 11.

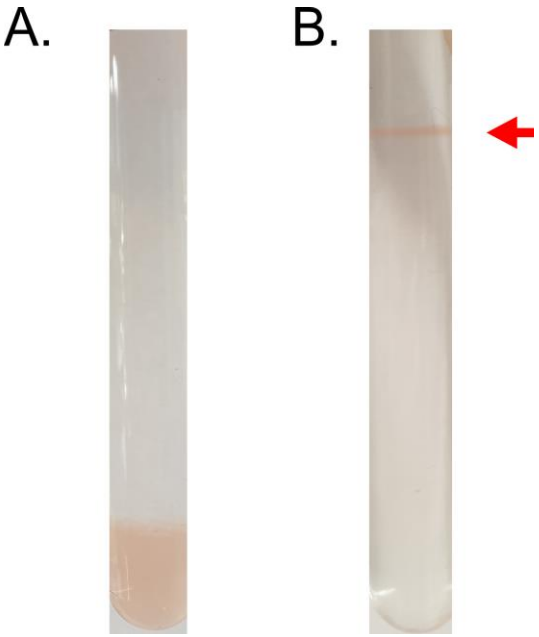


**Supplementary Figure 12. Optiprep™ density gradient of OapA knockout strain.** EVs **(A)** before and **(B)** after ultracentrifugation. Red arrow indicates where particles concentrated. For comparison with parental strain, see Supplementary Figure 1A.

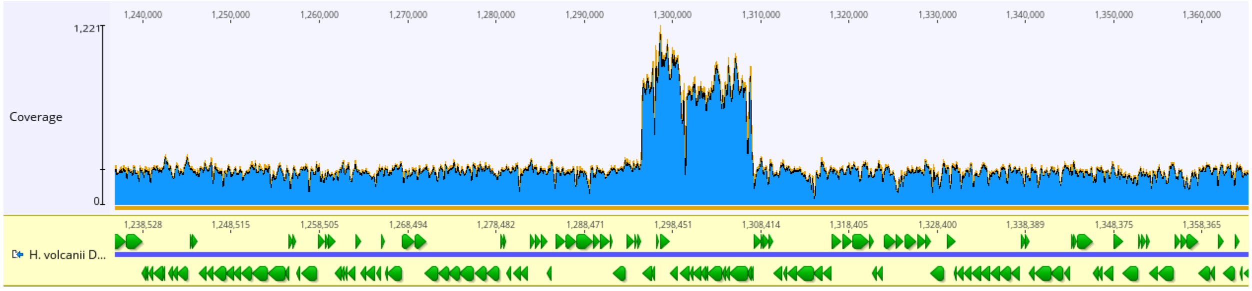


**Supplementary Figure 13. Resequencing of OapA knockout strain shows activation of proviral region.** Coverage blot of the genomic region including the provirus region. Image created in Geneious™.

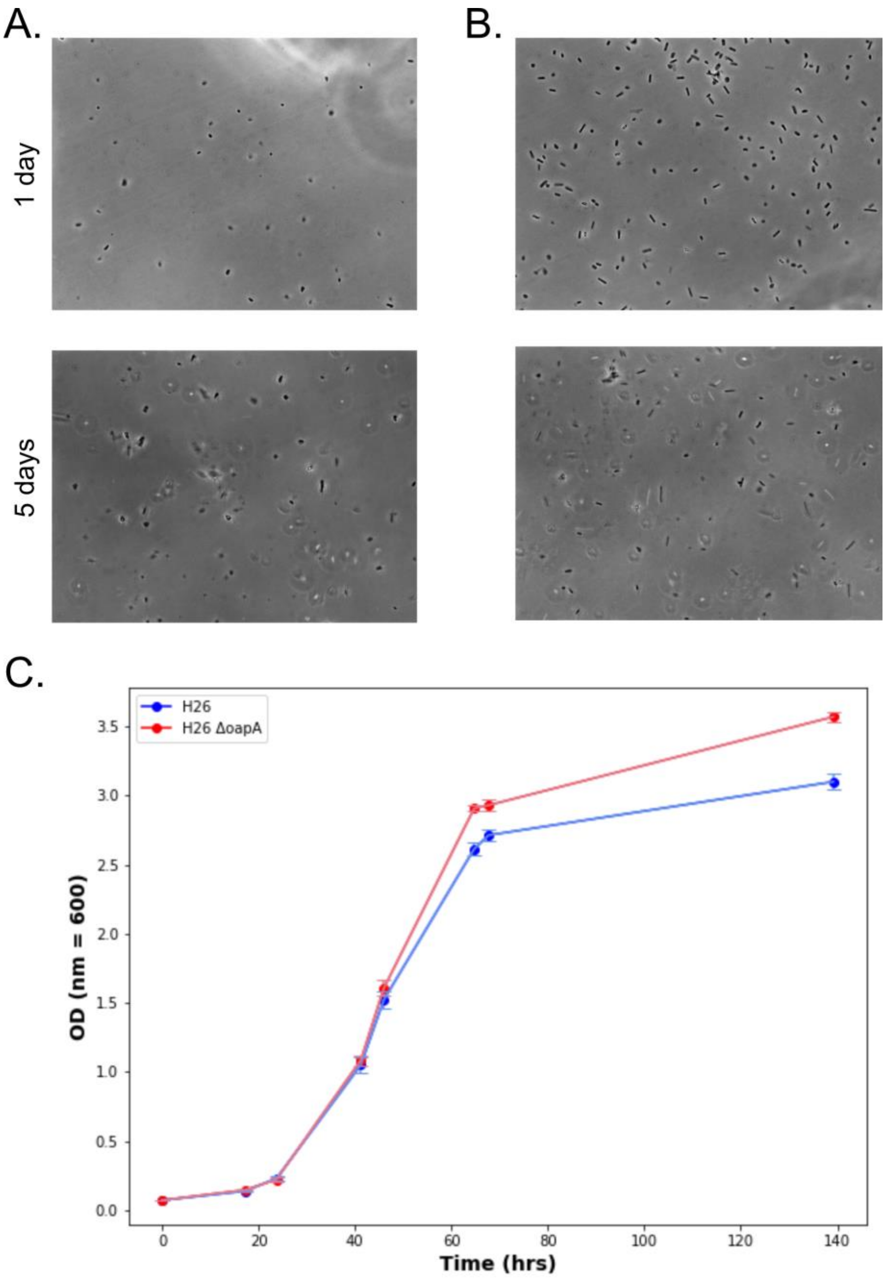


**Supplementary Figure 14. Phenotypes of OapA knockout strain in comparison to parental strain.** Phase contrast microscopy images of cells from OapA knockout strain **(A)** and parental strain **(B)** after 1 day (top) and 5 days (bottom) of growth. Samples were fixed with 1% glutaraldehyde and visualized with Axiophot Zeiss microscope. **(C)** Growth curve of parental strain (H26) and OapA knockout strain (H26 ΔoapA). Error bars represent standard deviation from three replicates.

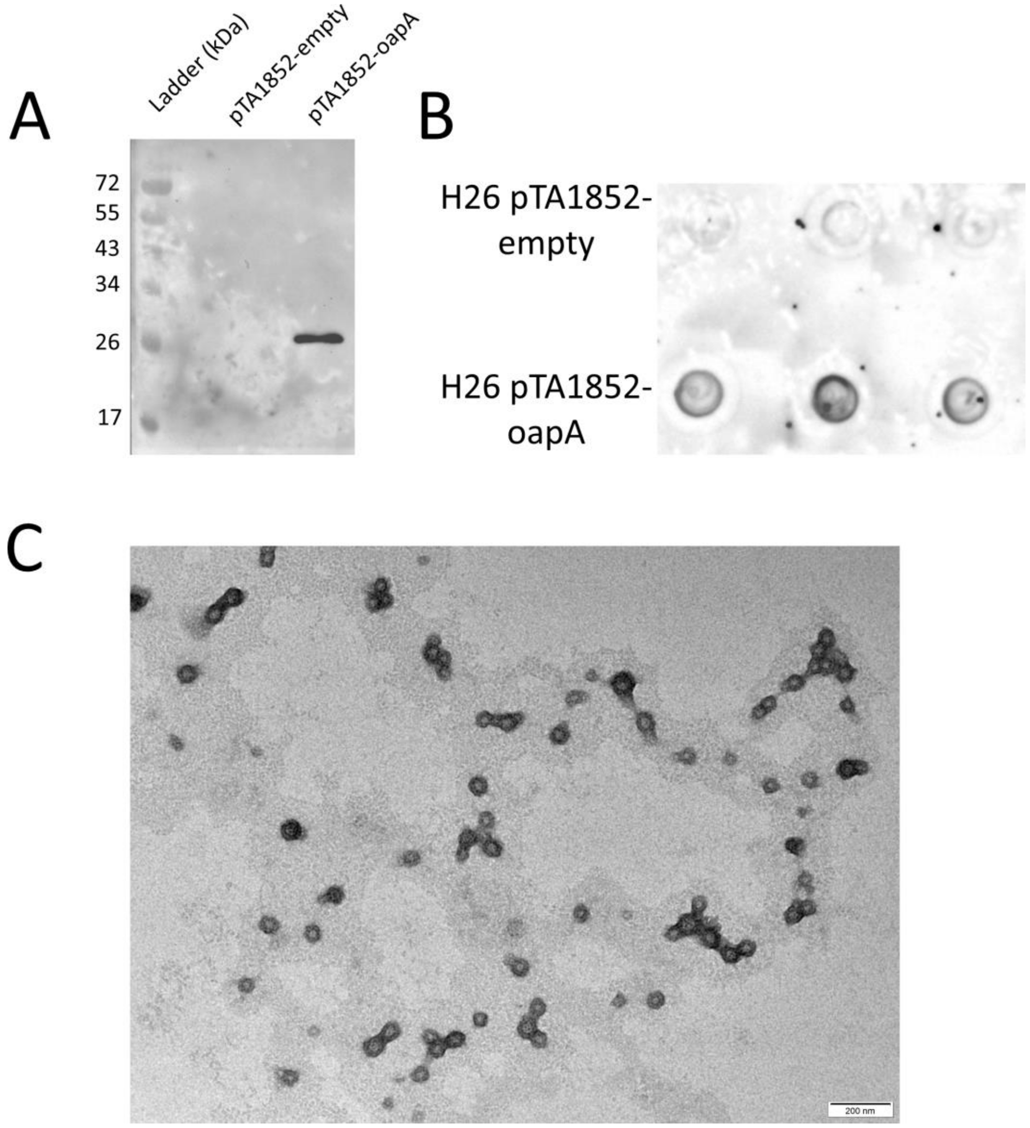


**Supplementary Figure 15. EV production in strains overexpressing OapA. (A)** Western blot with an anti-Strep tag antibody on affinity purified OapA expressed in H26, compared to an affinity purification from H26 with the empty vector (see methods). **(B)** Spot blot for quantification of EVs from cultures overexpressing OapA (Figure 5B). **(B)** Transmission electron micrographs of EVs isolated from strains overexpressing OapA. Scale bars denote 200 nm.

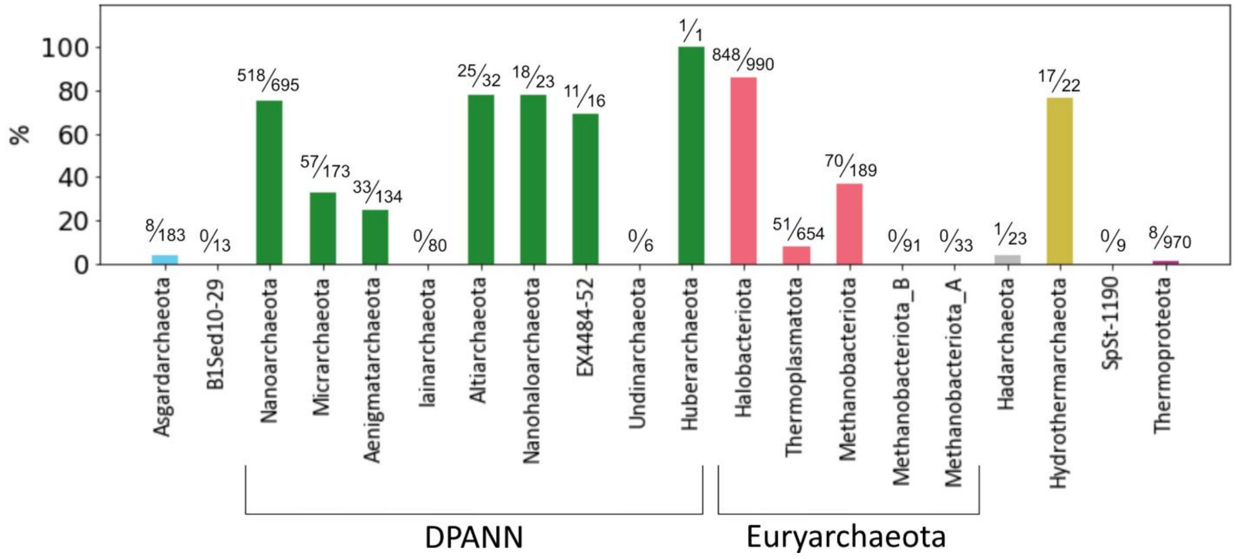


**Supplementary Figure 16. Archaeal Ras-family GTPase homologs across the archaeal domain.** Percentage of species identified containing a homolog for OapA within each phylum. Fraction above each bar denotes the number of species identified over the number of species surveyed.

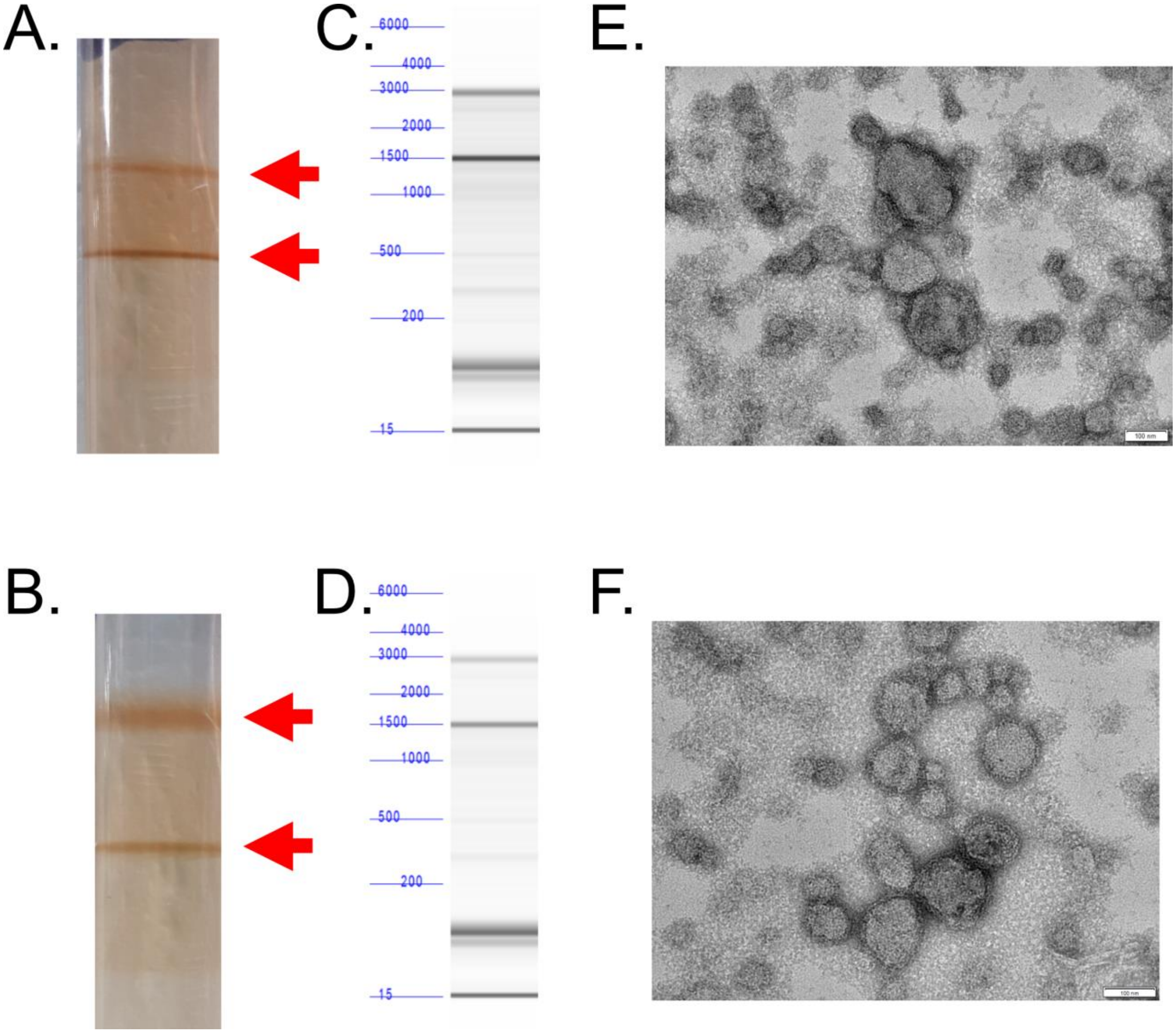


**Supplementary Figure 17. EVs isolated from CetZ1 and CetZ2 knockout strains.** Optiprep™ density gradients of EVs isolated from CetZ1 **(A)** and CetZ2 **(B)** knock out strains after ultracentrifugation. Red arrows indicate the upper and lower bands where particles had concentrated. **(C and D)** RNA was isolated from both strains and run on a fragment analyzer to observe the size distribution. For comparison to wild type, see Figure 2. **(E and F)** EVs could also be observed through TEM. Scale bars: 100 nm.

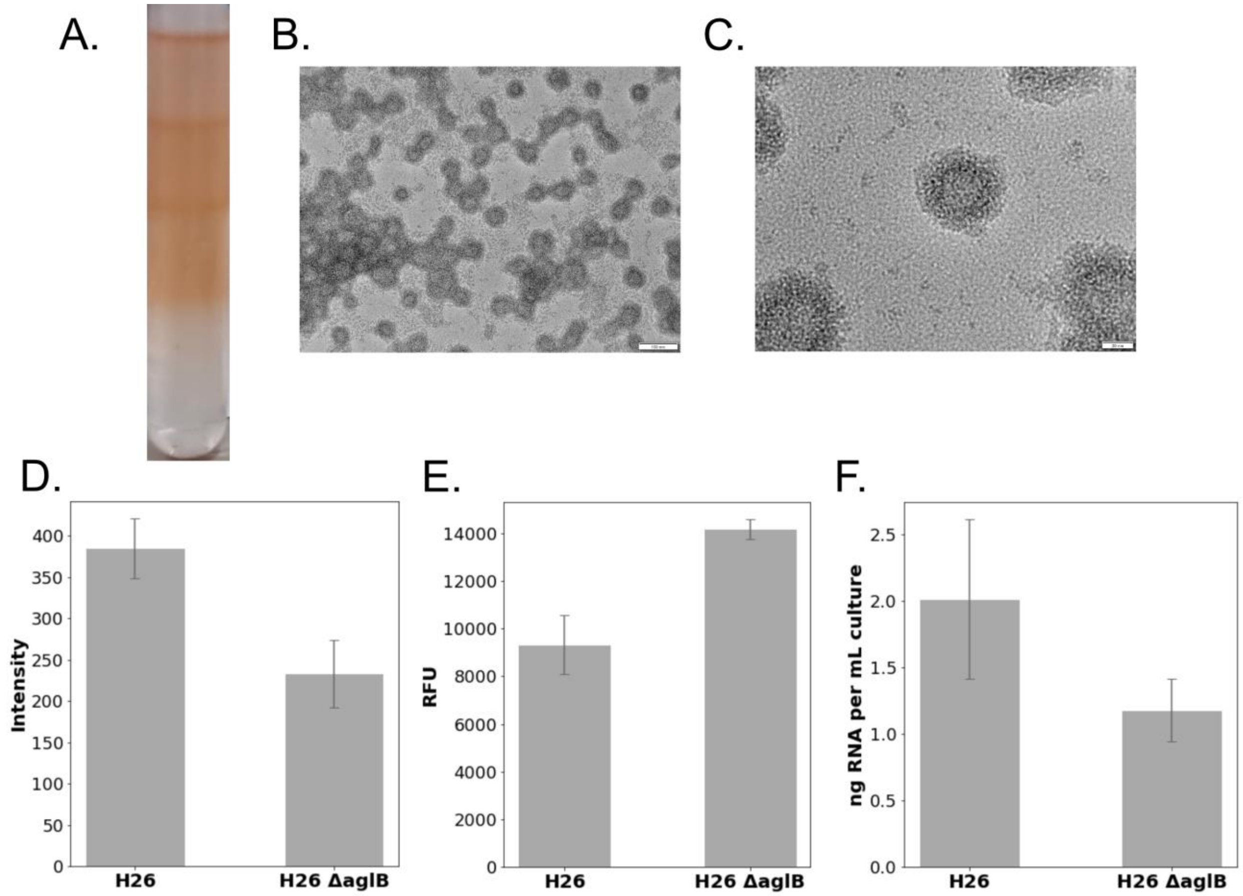


**Supplementary Figure 18. Phenotypes of EVs from AglB knockout strain. (A)** Bands of concentrated EVs after ultracentrifugation in Optiprep™ density gradient. Transmission electron microscopy of EVs isolated from AglB knockout strain with size bar 100 nm **(B)** and 20 nm **(C)**. Quantification of EV production of AglB knockout strain in comparison with parental strain by immunodetection **(D)** and fluorescence staining **(E)**. **(F)** Quantification of EV-associated RNA in culture supernatants of AglB knockout strain compared to parental strain normalized to OD (nm = 600). Error bars represent the standard deviation of three replicates. RFU = relative fluorescence unit.

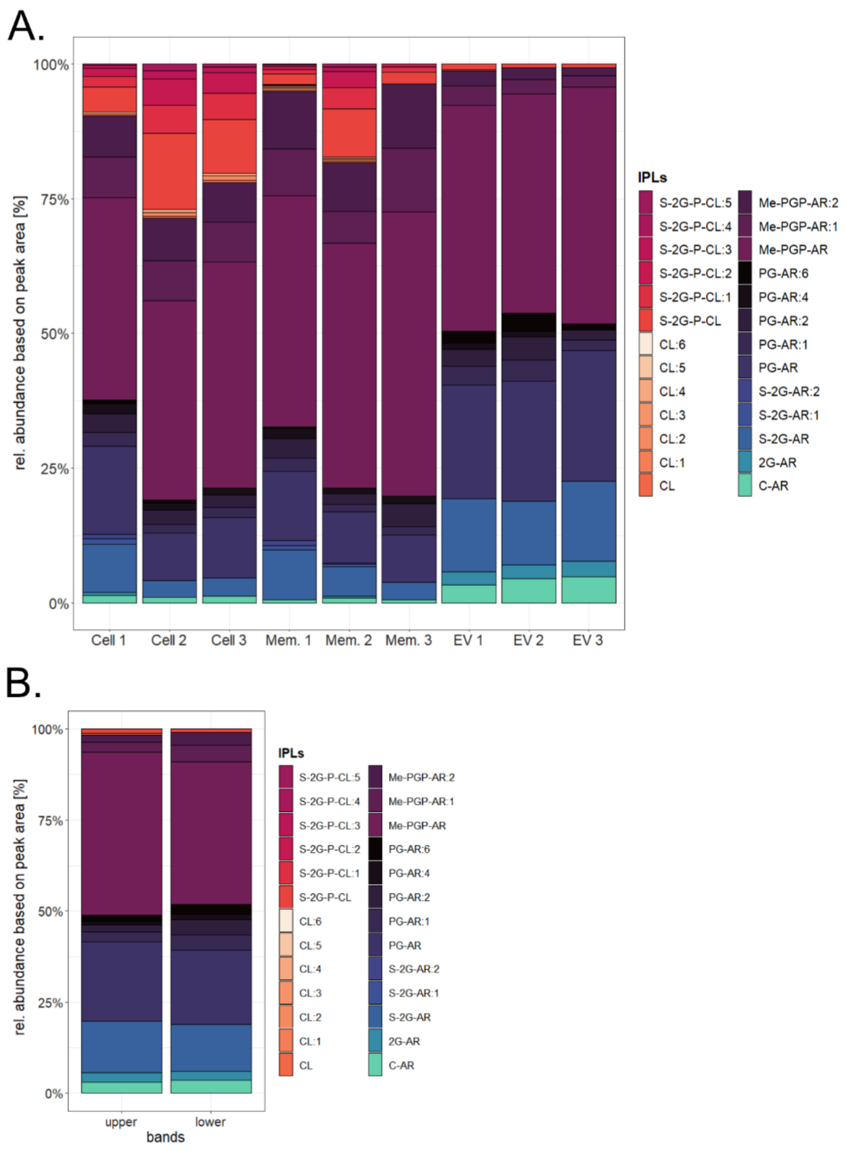


**Supplementary Figure 19. Distribution of lipid compounds comparing whole cells, cell membranes and EVs of H. volcanii. (A)** Individual replicates used to calculate the average relative abundances in Figure 6. Cell 1-3: whole cells, Mem. 1-3: membrane fraction and EV 1-3: extracellular vesicles after ultracentrifugation in Optiprep™ density gradients and bands pooled together for each biological replicate. **(B)** The lipid distribution in the upper (left column) and lower band (right column) after ultracentrifugation from one biological replicate. Relative abundances were calculated based on the peak area of the most abundant adduct for each compound. Lipids were identified based on their retention time, fractionation pattern and exact mass.

*Compound abbreviations*: AR = archaeol (C20-C20 isoprenoidal chains), CL = cardiolipin, :nUS = lipid with n number of unsaturations, UK = unknown compound. Lipids with neutral headgroups: 1G = monoglycosyl, 2G = diglycosyl, C-AR = core-AR. Lipids with anionic headgroups: PGP-Me = phosphatidylglycerophosphate methyl esters, PG = phosphatidylglycerol, S-2G =sulfated diglycosyl, S-GP = sulfoglycophospho, 2-PGLY = diphosphoglycerol.

### Supplementary Tables

#### Supplementary Table 1: Strains used in this study

| Name | Reference | Organism | Description | Media | Supplement |
|---|---|---|---|---|---|
| *H. volcanii DS2* | [6] | *H. volcanii DS2* | Wild type strain | HV-cab/YPC | none |
| H26 | [7] | *H. volcanii* | ΔpyrE2 | HV-cab/YPC | uracil |
| H26 ΔoapA | [8] | *H. volcanii* | ΔpyrE2, ΔoapA | HV-cab/YPC | uracil |
| H26 ΔaglB | [9] | *H. volcanii* | ΔpyrE2, ΔaglB | HV-cab/YPC | uracil |
| H53 | [7] | *H. volcanii* | ΔpyrE2, ΔtrpA | HV-cab/YPC | uracil, tryptophan |
| H53 ΔcetZ1 | [10] | *H. volcanii* | ΔpyrE2, ΔtrpA, ΔcetZ1 | HV-cab/YPC | uracil, tryptophan |
| H53 ΔcetZ2 | [10] | *H. volcanii* | ΔpyrE2, ΔtrpA, ΔcetZ2 | HV-cab/YPC | uracil, tryptophan |
| *Halobacterium salinarum* | [11] | *Halobacterium salinarum* | Wild type strain | HS-Media | none |
| *Halorubrum lacusprofundi DL18* | [12] | *Halorubrum lacusprofundi DL18* | Wild type strain | DBCM2 | none |

#### Supplementary Table 2: Primer Sequences

| Name | Oligonucleotide sequence 5'-3' | Description |
|---|---|---|
| HFPV1F | CACGAACGAGAACACCGACC | Forward primer to test infection of HFPV-1 |
| HFPV1R | TGATGACGAATCCAACGAGCAG | Reverse primer to test infection of HFPV-1 |
| AglB_US_F | CCGGCCAAGCTTGGTTTGCGAGCGACCCAGTCG | Forward primer for upstream flank of aglB with HindIII restriction sites |
| AglB_US_R | GAATTCGCCGCCCGAAGATCTTGTGACCAACAACCGCCAAG | Reverse primer for upstream flank of aglB with EcoRI and BglII restriction sites |
| AglB_DS_F | AGATCTTCGGGCGGCGAATTCCACGAGCCGAGACGGCGACGA | Forward primer of downstream flank of aglB with BglII and EcoRI restriction sites |
| AglB_DS_R | CCGGCCGGATCCGCGCGTCGCCGTGCTCGGAC | Reverse primer for downstream flank of AglB with BamHI restriction sites |
| csg probe | GCTGTCAGCGTCGAGGTTTCC | Northern blot probe for the 5' end of S-layer mRNA |
